# *oskar* acts with the transcription factor Creb to regulate long-term memory in crickets

**DOI:** 10.1101/2022.10.24.513429

**Authors:** Arpita Kulkarni, Ben Ewen-Campen, Kanta Terao, Yukihisa Matsumoto, Yaolong Li, Takayuki Watanabe, Jonchee A. Kao, Swapnil S. Parhad, Guillem Ylla, Makoto Mizunami, Cassandra G. Extavour

## Abstract

Novel genes have the potential to drive the evolution of new biological mechanisms, or to integrate into pre-existing regulatory circuits and contribute to the regulation of older, conserved biological functions. One such gene, the novel insect-specific gene *oskar*, was first identified based on its role in establishing the *Drosophila melanogaster* germ line. We previously showed that this gene likely arose through an unusual domain transfer event involving bacterial endosymbionts, and played a somatic role before evolving its well-known germ line function. Here, we provide empirical support for this hypothesis in the form of evidence for a novel neural role for *oskar*. We show that *oskar* is expressed in the adult neural stem cells of a hemimetabolous insect, the cricket *Gryllus bimaculatus*. In these stem cells, called neuroblasts, *oskar* is required together with the ancient animal transcription factor *Creb* to regulate long-term (but not short-term) olfactory memory. We provide evidence that *oskar* positively regulates *Creb*, which plays a conserved role in long-term memory across animals, and that *oskar* in turn may be a direct target of Creb. Together with previous reports of a role for *oskar* in nervous system development and function in crickets and flies, our results are consistent with the hypothesis that *oskar*’s original somatic role may have been in the insect nervous system. Moreover, its co-localization and functional cooperation with the conserved pluripotency gene *piwi* in the nervous system may have facilitated *oskar*’s later co-option to the germ line in holometabolous insects.

## Introduction

*Oskar* (*osk*) is an insect-specific gene first discovered in *Drosophila melanogaster*, where it plays a critical role in germline specification (1). Oskar mRNA is localized to the posterior of the developing *D. melanogaster* oocyte (2, 3). Local translation and anchoring of Oskar (Osk) protein leads to the posterior accumulation of the mRNA and protein products of several genes with conserved expression and function in animal germ lines, including *vasa* and *piwi* (2, 4, 5). Collectively called germ plasm, these cytoplasmic contents act as necessary and sufficient determinants to specify embryonic germ cells (2, 3). The current model of Osk function in *D. melanogaster* germ plasm assembly is that it serves as a scaffolding protein, facilitating the assembly of the ribonucleoprotein complexes that contain germ plasm components (2, 6, 7).

Interestingly, *osk* and several other genes originally identified as *D. melanogaster* germ line genes, including *vasa*, *pumilio*, *staufen*, *orb*, and *piwi*-related genes including *aubergine* and *argonaute 3*, have since been shown to have a variety of roles in animal nervous systems(8–14). For example, in *D. melanogaster*, *osk* RNAi in larval dendritic arborization neurons disrupts *nanos* mRNA localization, ultimately leading to a defect in dendrite morphogenesis and an associated defect in motor response to mechanical stimulation(12). Furthermore, *osk* plays a role in the embryonic nervous system, but not in the germ line, in a hemimetabolous insect, the cricket *Gryllus bimaculatus*, where it is important for proper neuroblast divisions and subsequent axonal patterning(15). Our recent analysis of hundreds of previously unidentified *osk* orthologs across insects showed that *osk* is expressed in at least a dozen somatic tissues in species across the insect tree(16). This suggests that a somatic function of *osk* may be ancestral. However, the precise roles of *osk* in any somatic tissue, including the nervous system, remain largely unknown.

Here, we demonstrate a role for *osk* in the adult brain of the cricket *Gryllus bimaculatus*, in a population of neural stem cells in the mushroom body that persist throughout adult life. We show that *osk*, as well as Piwi and Vasa, are enriched in a population of adult neuroblasts in the mushroom body, and that RNA interference (RNAi) targeting *osk* or *piwi* in adult crickets impairs long-term, but not short-term memory formation in an olfactory associative learning assay(17). We also provide evidence that *osk* and *piwi* function in a regulatory feedback loop with the <u>c</u>yclic AMP <u>r</u>esponse <u>e</u>lement <u>b</u>inding protein (Creb), a transcription factor with well described conserved roles in long-term memory across metazoans(18). Our data demonstrate a novel somatic role for *osk*, and shed light on how a novel gene may acquire critical roles by integrating with pre-existing gene regulatory systems comprising older, conserved genes.

## Results

### osk is expressed in adult neuroblasts of the mushroom body

We previously showed that neuroblasts in the cricket embryo express *osk*, *vasa* and *piwi*, and that *osk* is required for correct neuroblast division and embryonic nervous system morphology(15). Interestingly, in many insects, including crickets, a subset of embryonic neuroblasts persist in the brain throughout adulthood and continuously give rise to new neurons called Kenyon cells that comprise the mushroom body(19–21). This contrasts with flies like *D. melanogaster*, in which neuroblasts die prior to adulthood(22), and in which adult brains are thus essentially devoid of neurogenesis(23) (although there are reports of potential stem cells in adult *D. melanogaster* brains(23, 24) which may be damage-dependent rather than homeostatic in function(25), and which remain controversial (26)).

Given the role of *osk* in embryonic neuroblasts of crickets, we asked whether *osk* also plays a role in the adult mushroom body neuroblasts. We used *in situ* hybridization to examine *osk* expression in the adult brain, and found expression in a cluster of cells with the large, round nuclei and diffuse chromatin characteristic of stem cells, at the apex of each of the two lobes of the mushroom body, consistent with descriptions of adult neuroblasts in orthopterans (Fig. 1A). EdU co-localization (Fig.1B) confirmed the identity of these cells as neuroblasts, the only proliferative cells in the adult brain(27). We also found that mushroom body neuroblasts express high levels of Vasa and Piwi proteins (Fig. 1E).

**Figure 1.**
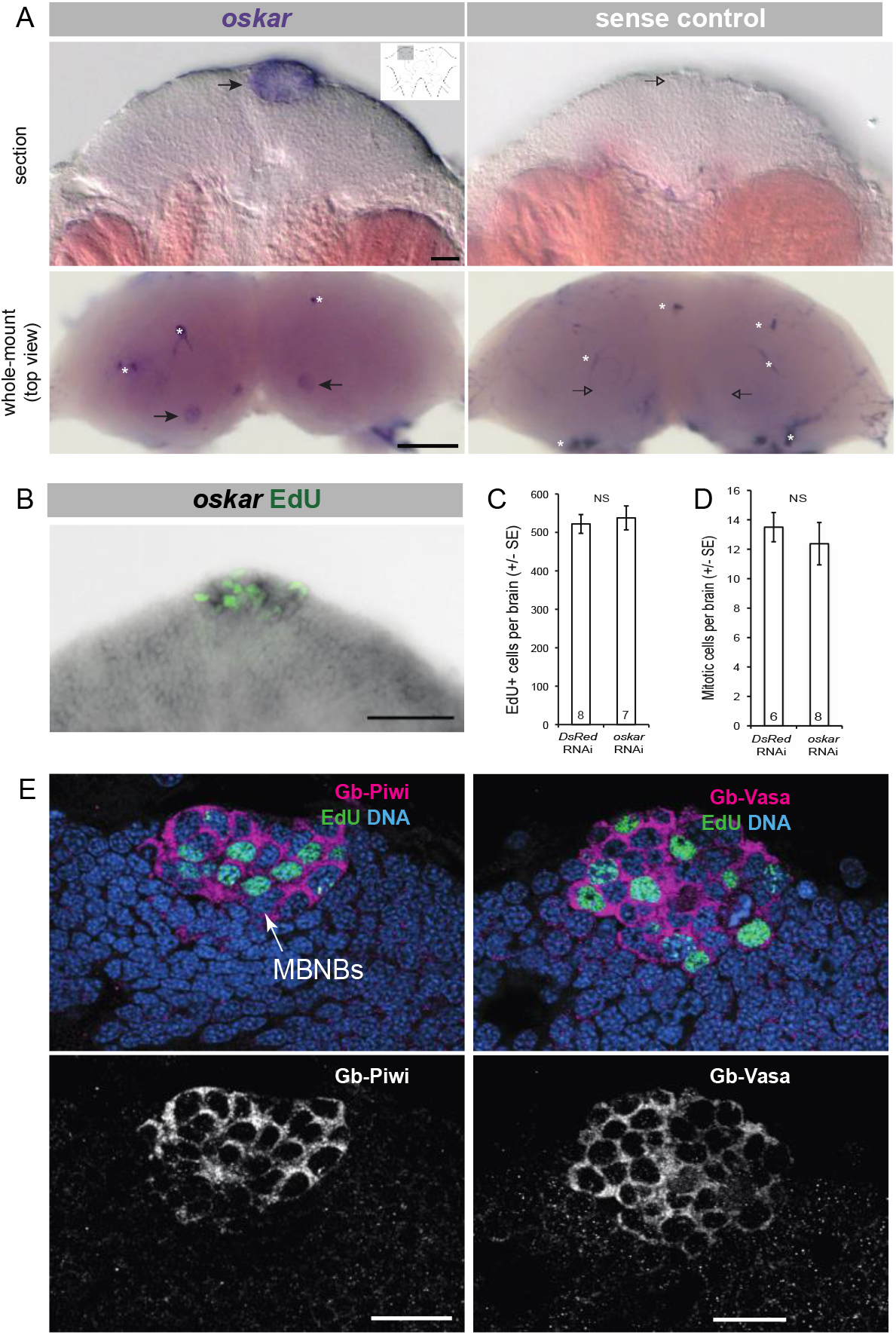
*oskar*, Piwi and Vasa are expressed in *G. bimaculatus* adult mushroom body neuroblasts. (**A**) In situ hybridization on adult *G. bimaculatus* brains detects *osk* transcripts in the cells of the mushroom body (arrows). Inset in top left panel shows the overall structure of the adult brain; shaded box indicates a single mushroom body lobe, corresponding to the region shown in micrographs in top row. Bottom row: dorsal views of both mushroom body lobes, indicating *oskar* expression in neuroblast clusters (arrows). White asterisks indicate non-specific binding of probe to tracheal remnants in the brain. (**B**) Edu labelling (green) of the adult brain shows that *osk*-expressing cells (grey) are mitotically active, consistent with their identity as neuroblasts. **(C)** Quantification of EdU-positive cells shows no significant difference between *osk^RNAi^* and control brains (p<0.05). **(D)** Quantification of total number of mitotically active cells shows no significant difference between *osk^RNAi^* and control brains (p<0.05). Numbers within bars indicate sample sizes in (**C**) and (**D**). **(E)** Detection of Vasa & Piwi proteins (magenta) in adult mushroom body neuroblasts (MBNBs). Scale bars = 50μm in top panels of (**A)** and (**B)** and in **(E)**, and 200μM in bottom panels of (**A)**.

Previous research has shown that mushroom body neuroblasts play an important role in long-term olfactory memory formation in Orthoptera(28). Scotto-Lomassesse(28) found that ablation of mushroom body neuroblasts using irradiation led to a dramatic reduction in olfactory, but not visual, learning after 24- and 48-hours, suggesting that newborn mushroom body neurons produced by those neuroblasts play a role in forming new olfactory memories. We therefore sought to test whether *osk*, expressed specifically in mushroom body neuroblasts, functions in these cells in the context of long-term memory formation.

We first tested whether Osk regulates the proliferation or survival of adult mushroom body neuroblasts. Using an established technique for systemic RNAi in the adult cricket brain(29), we injected double stranded *osk* RNA (dsRNA) into the head capsule, and confirmed efficiency of *osk* knockdown via qPCR (Fig. 2D; Suppl. Table S1) and small RNA profiling of *osk^RNAi^* brains (Suppl. Tables S2, S3 and S4; Suppl. Fig. S1). *osk^RNAi^* adult brains showed no gross anatomical defects relative to controls (data not shown). Moreover, neither the total number of neuroblasts (p<0.05), nor the number of neuroblasts undergoing mitosis as revealed by EdU labeling (p<0.05), were statistically significantly different between *osk^RNAi^* adult brains and controls (Fig. 1C, 1D). We stained *osk^RNAi^* and control brains with Cleaved Caspase-3, a marker for apoptosis, and did not observe any evidence of cell death (Suppl. Fig. S2A). We noted that one described role for *piwi* in the *Drosophila* germ line is to prevent DNA damage caused by transposon mobilization(30). However, we observed no detectable increase in γH2A staining, a marker for DNA damage, in *osk^RNAi^* brains (Suppl. Fig. S2B). These data suggest that *osk* is not required for the proliferation, survival or genomic integrity of adult neuroblasts. However, the specific expression of *osk*, Piwi, and Vasa in the mushroom body neuroblasts suggested that some or all of these genes could play a role related to memory or learning.

**Figure 2.**
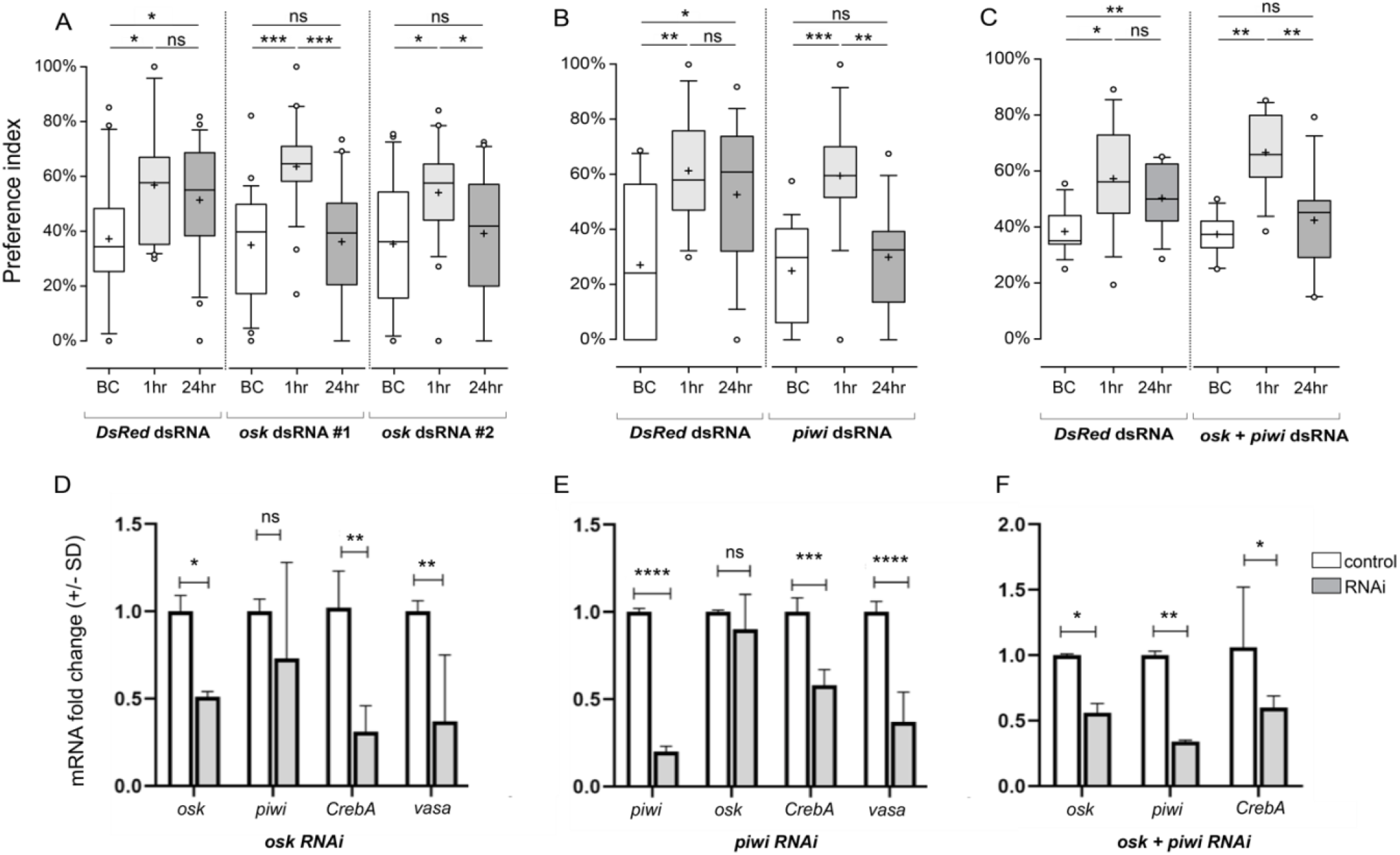
*oskar^RNAi^* and *piwi^RNAi^* impairs cricket long-term memory. (**A**) Results of olfactory memory assay in *osk^RNAi^* animals. (**B**) Results of olfactory memory assay in *piwi^RNAi^* animals. (**C**) Results of olfactory memory assay in *osk^RNAi^/piwi^RNAi^* double knockdown animals. For each assay, relative preference between the rewarded odor (peppermint) and control odor (vanilla) was tested before conditioning (BC), one hour post-training (1hr), and one day post training (24hr) for *DsRed^RNAi^* controls and for *osk^RNAi^* (using two different non-overlapping *osk* fragments #1 and #2), *piwi^RNAi^* and *osk^RNAi^/piwi^RNAi^*. Boxes represent the 1^st^ and 3^rd^ quartiles surrounding the median (middle line). Whiskers extend to values within 1.5x of interquartile range. Wilcoxon’s test was used for comparison of preference before and after conditioning. For multiple comparisons, the Holm method was used to adjust the significance level (* p < 0.05, ** p<0.01, *** p<0.001, n.s. = not statistically significant). (**D-F)**. qPCR results showing the extent of downregulation of different *G. bimaculatus* genes in *osk^RNAi^, piwi^RNAi^* and *osk^RNAi^/piwi^RNAi^* backgrounds. Effectiveness of RNAi per background is also shown in each case. Data is plotted as mRNA fold change (+/- standard deviation) based on the ΔΔCt method (* p < 0.05, ** p < 0.01, *** p<0.001, **** p < 0.0001, n.s. = not statistically significant).

### oskar RNAi impairs long-term but not short-term memory

The mushroom body is the anatomical substrate for olfactory memory and learning in insects(27, 31, 32), and ablation of the mushroom body or of the adult mushroom body neuroblasts impairs these processes(28, 33, 34). Based on previous observations that mushroom body neuroblasts play a role in long-term olfactory memory formation in crickets(28), we hypothesized that *osk* might play a role in this process. To test this hypothesis, we assessed memory of *osk^RNAi^* adult male crickets at one hour (“short-term memory”) and one day (“long-term memory”) post-training using well-established cricket olfactory behavior assays as previously described (17). Briefly, crickets were injected with double stranded RNA against the gene of interest (*oskar*) or a control gene (*DsRed* (35), and then subjected to odor preference tests (allowed to freely visit peppermint and vanilla odor sources, quantifying the time spent at each odor source) and conditioning trials (peppermint odor was paired with a water reward, and relative preference for this rewarded odor was compared before and after the conditioning) (17).

In control crickets (injected with dsRNA targeting *DsRed*(35)), four training sessions led to a significant (p<0.05) short-term preference for the rewarded odor (peppermint) at one hour after training (short-term; Fig. 2A). Trained control crickets retained this learned preference (p<0.01) even at one day after training (Fig. 2A, “*DsRed*”), demonstrating that long-term memory formation is intact in these controls. However, although *osk^RNAi^* crickets formed and retained memory for the rewarded odor by one hour after training (short-term; Fig.2A *osk* dsRNA #1, p<0.001) this memory was lost by one day after training (long-term; Fig. 2A *osk* dsRNA #1, p>0.05), indicating a specific impairment of long-term memory formation. These results were reproducible in a second experiment using a non-overlapping fragment of *osk* dsRNA (Fig. 2A, *osk* dsRNA #2; p<0.05 for short-term, p>0.05 for long-term), suggesting the impact was specific to *osk* knockdown. Efficacy of the knockdown was confirmed via qPCR (Fig. 2D; Suppl. Table S1) and small RNA sequencing (Suppl. Fig. S1, Suppl. Tables S1, S2), indicating that *osk* is required for cricket long-term memory.

Since both Piwi and Vasa were co-expressed with *osk* in cricket mushroom body neuroblasts(15) (Fig. 1E), we also assessed the role of these two genes in olfactory memory. We found that *piwi* (Fig. 2B, p<0.001) but not *vasa* (Suppl. Fig. S3A, p>0.05; Suppl. Fig. S3B) was also required for cricket long-term memory. qPCR analyses showed that *osk* RNAi led to a significant decrease in *piwi* transcript levels (Fig. 2D), suggesting that *osk* positively regulates *piwi* in the cricket brain. However, *osk* transcript levels remained unaffected in *piwi^RNAi^* animals (Fig. 2E). Consistent with the phenotype of the single gene knockdowns, *osk^RNAi^/piwi^RNAi^* double knockdown adults also showed a long-term memory impairment phenotype (Fig. 2C, p<0.01). Thus, *osk* and *piwi* do not globally disrupt olfaction, learning or short-term memory formation, but are required for consolidation of long-term memory in this species.

### Osk and piwi positively regulate the nuclear transcription factor CrebA

To understand how a novel gene like *osk* might have gained a role in an ancient animal function like long-term memory consolidation, we investigated the hypothesis that it might interact with conserved regulators of animal memory. To test this hypothesis, we asked whether we could detect a functional or regulatory interaction between *osk* and a highly conserved transcription factor with well-documented roles in long-term memory formation across animals, <u>c</u>yclic AMP <u>r</u>esponse <u>e</u>lement <u>b</u>inding protein (Creb)(18). We first identified putative *Creb* orthologs in the *G. bimaculatus* genome(36) using a combination of BLAST searches and phylogenetic analyses (Fig. 3A, Suppl. Table S5). These analyses yielded two high-confidence *Creb* orthologs, which we called *CrebA* and *CrebB* based on their closest *D. melanogaster Creb* gene relative (Fig. 3A). Analysis of previously generated transcriptomes(37) showed that both genes are expressed in adult cricket brains (Suppl. Table S6). We performed *CrebA* and *CrebB* RNAi experiments and discovered that *CrebA* (but not *CrebB;* Suppl. Fig. S3C) was required for long-term memory in crickets (Fig. 3B; p<0.003 and p<0.001 for dsRNA#1 and dsRNA#2 respectively; Suppl. Fig. S3C). Using quantitative PCR (qPCR), we then asked whether transcript levels of this memory regulator were altered in *osk* or *piwi* knockdown conditions, and found consistent downregulation of *CrebA* transcript levels in both single and double RNAi backgrounds (Fig. 2D-F). In contrast, and consistent with the observation that *vasa^RNAi^* had no long-term memory impact (Suppl. Fig. 3A), qPCR revealed no reduction of *CrebA* transcripts in *vasa^RNAi^* conditions (Suppl. Fig. 3B). This suggests that the long-term memory defects observed in *osk^RNAi^* and *piwi^RNAi^* conditions (Fig. 2D-F) are due to a downregulation of *CrebA* in these animals.

**Figure 3.**
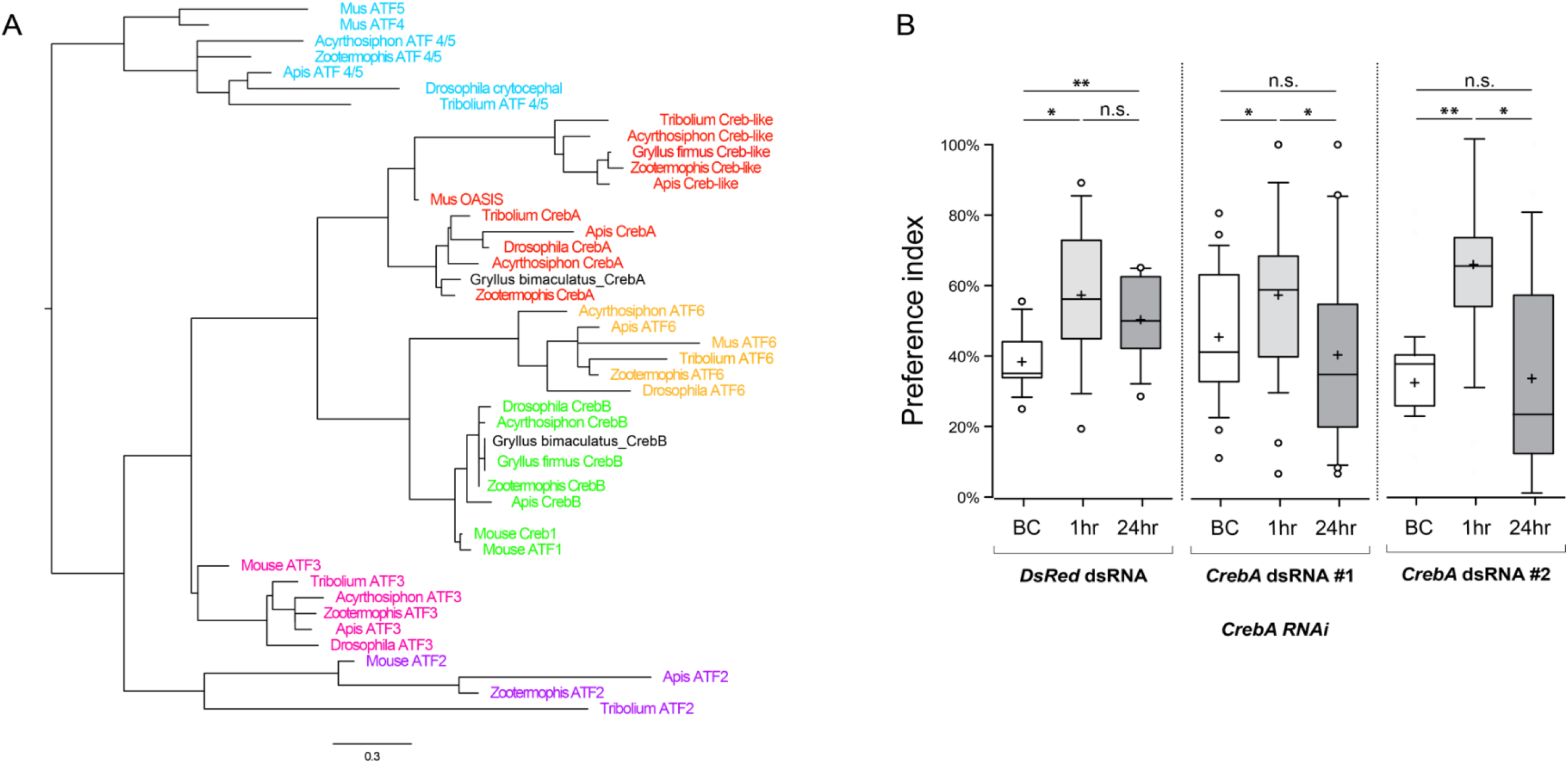
Cricket *CrebA* is required for cricket long-term memory. (**A**) GenBank IDs of *Creb/ATF* family member orthologs in mouse and insects (Suppl. Table S5) were used to construct a *Creb* phylogenetic tree to infer the evolutionary relationships between mammalian *Creb* proteins and their insect counterparts. *G. bimaculatus CrebA* and *CrebB* are indicated in black in the tree (**B**) *CrebA*^RNAi^ impairs long-term memory formation in crickets. Relative preference between the rewarded odor (peppermint) and control odor (vanilla) was tested before conditioning (BC), one hour post-training (1hr), and one day post training (24hr) for *DsRed^RNAi^* controls and *CrebA^RNAi^* (using two different non-overlapping *CrebA* fragments #1 and #2 for independent confirmation). Boxes represent the 1^st^ and 3^rd^ quartiles surrounding the median (middle line). Whiskers extend to values within 1.5x of interquartile range. Wilcoxon’s test was used for comparison of preference before and after conditioning. For multiple comparisons, the Holm method was used to adjust the significance level. (* p < 0.05, ** p<0.01, *** p<0.001, n.s. = not statistically significant). n=9 for *CrebA* and n=10 for *DsRed*.

### Osk and piwi are regulated by CrebA

Creb proteins are transcription factors that bind <u>c</u>yclic AMP <u>r</u>esponse <u>e</u>lement (CRE) binding sites within the regulatory regions of target genes to initiate transcription(18) (Fig. 4A). Since target gene transcription and new protein synthesis is crucial for long-term memory formation, and given the similarity in long-term memory phenotypes of *osk^RNAi^*, *piwi^RNAi^* and *CrebA^RNAi^* animals, we asked whether *osk* or *piwi* might also be *Creb* target genes in this cricket. qPCR revealed that transcript levels of both *osk* and *piwi* are significantly decreased in *CrebA^RNAi^* conditions (Fig. 4B), suggesting that *osk/piwi* and CrebA may interact in a positive feedback loop to regulate each other’s transcript levels. To evaluate the possibility that *osk* or *piwi* might be direct transcriptional targets of CrebA, we examined the genomic sequences within 10kb upstream of both loci and found two bioinformatically predicted CRE binding sites within the 6kb upstream of the transcription start sites for *osk* and *piwi* (Fig. 4C; Suppl. File 1). These predicted CRE binding sites were found twice as frequently as we would expect to find such sequences in a randomly generated sequence of this length (see Methods). Electrophoretic mobility shift assays showed that protein(s) within the adult cricket brain bind specifically to the predicted CRE sites of *osk* (Fig. 4D; Suppl. Table S7). Given the current lack of species-specific CrebA reagents for this cricket species, we cannot rule out the interpretation that a protein(s) other than CrebA present in the adult cricket brain is causing the observed mobility shift by binding the predicted CRE sites of *osk*. However, given our functional data indicating that RNAi against *osk*, *piwi* and *CrebA* all yield long-term memory defects (Figs. 2, 3B), that *osk* and *piwi* transcript levels are reduced in *CrebA^RNAi^* brains (Fig. 4B), and that *osk* and *piwi* genomic loci contain predicted CRE binding sites (Fig. 4C), the results of our gel shift assay are consistent with the hypothesis that cricket *osk* is a direct transcriptional target of CrebA.

**Figure 4.**
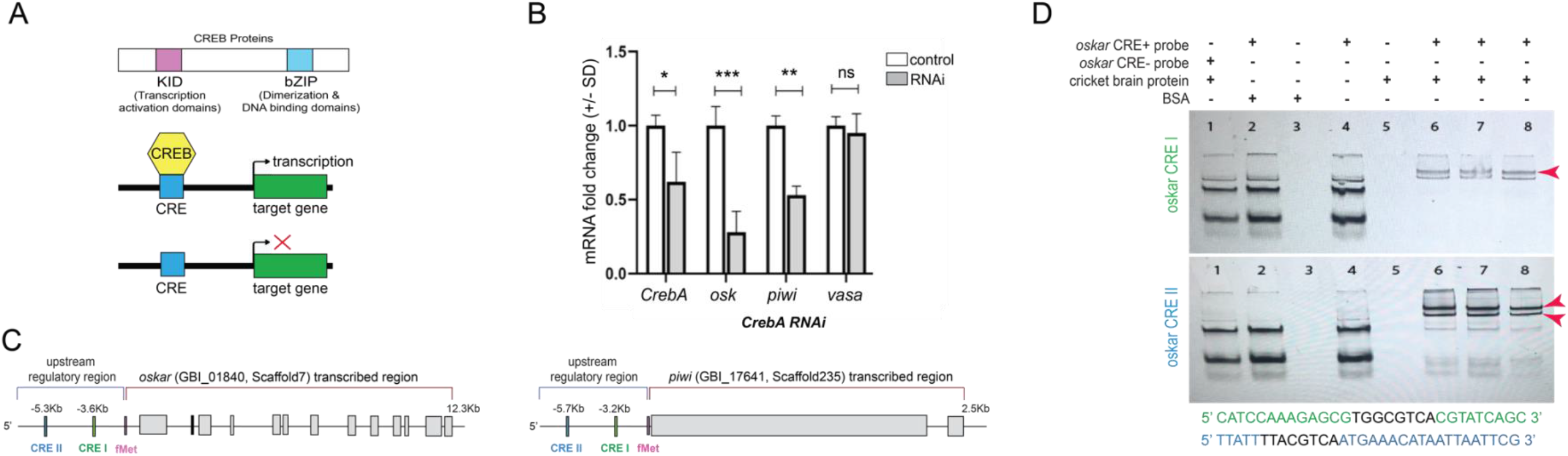
Cricket CrebA regulates *oskar*. (**A**) Schematic diagram of the transcription factor <u>c</u>AMP <u>re</u>sponse <u>e</u>lement <u>b</u>inding protein (Creb) protein (top) displaying only the two domains relevant to this study, the <u>K</u>inase-<u>I</u>nducible <u>D</u>omain (KID) that can facilitate kinase-inducible transcription activation and the <u>b</u>asic leucine <u>zip</u>per (bZIP) domain that is important for dimerization and DNA binding. Creb proteins bind the <u>c</u>yclic AMP <u>r</u>esponse <u>e</u>lement (CRE), a sequence present in the promoter regions of many cellular genes to increase (middle schematic) or decrease (bottom schematic) transcription of target genes. (**B**) qPCR results showing the relative expression levels of *G. bimaculatus osk*, *piwi*, and *vasa* in *CrebA^RNAi^* knockdown conditions. The extent of *CrebA* transcript decrease is also shown to assess efficiency of RNAi knockdown. Data is plotted as mRNA fold change (+/- standard deviation) based on the ΔΔCt method (* p < 0.05, ** p < 0.01, *** p<0.001, * p < 0.0001 n.s. = not statistically significant). (**C**) Schematic of *G. bimaculatus oskar* and *piwi* genes showing exons (depicted as rectangular grey boxes) along with their presumptive upstream regulatory regions, each containing two predicted CRE sites, which we call CRE I (~3.6Kb and ~3.2Kb upstream of the predicted transcription start site for *osk* and *piwi* respectively, marked by fMet) and CRE II (~5.3Kb and ~5.7Kb upstream of predicted fMet for *osk* and *piwi* respectively). (**D**) <u>E</u>lectrophoretic <u>M</u>obility <u>S</u>hift <u>A</u>ssay (EMSA) to detect possible Creb binding to *osk’s* CRE I (top, in green) and CRE II (bottom, in blue). “*oskar* CRE+ probe” indicates predicted CRE site-containing probe “*oskar* CRE-probe” indicates probe without predicted CRE sites; “cricket brain protein” indicates *G. bimaculatus* brain protein extract; “BSA” indicates 1% BSA as a non-specific protein control. The complete sequences of the EMSA probes used for CRE I and II experiments are indicated in green and blue text underneath the gel image, with black representing the predicted CRE sequence. Shift marked with red arrowheads.

## Discussion

We have discovered a new role for *oskar* in the adult cricket brain (Fig.1). We have shown that *osk*, Piwi, and Vasa are co-expressed in mushroom body neuroblasts (Fig. 1A, B), a population of stem neural cells required for long-term olfactory memory formation(28), and that knockdown of *osk* and *piwi* disrupts olfactory long-term memory formation. The precise role that the mushroom body neuroblasts play in memory formation remains unknown, as does the molecular role of Osk in these cells. In *D. melanogaster*, where there are no adult neural stem cells in the mushroom body, olfactory long-term memory requires the ~2,500 differentiated Kenyon cell neurons of the mushroom body (38), which respond with high selectivity to a small number of stimuli, allowing the mushroom body to house an explicit representation of a large number of olfactory cues(38, 39). Specific olfactory stimuli are associated with learned behavioral responses via specific sets of neurons connecting the mushroom body to other brain regions in a protein synthesis-dependent fashion, to form long-term memories(9, 40). Thus, one possibility is that adult-born Kenyon cells in *G. bimaculatus* (and other insects that display adult neurogenesis in the mushroom body) are recruited into an existing circuit and allow for a constantly increasing repertoire of olfactory associations. Our results suggest that *osk* could play a role in this process, as *osk* RNAi disrupts long-term memory. We note that of the two mammalian brain regions known to undergo adult neurogenesis, one (the subventricular zone) contributes to the olfactory bulb, and neurogenesis in this region is involved in olfactory memory (41).

Given that adult *D. melanogaster* lacks the mushroom body neuroblasts seen in *G. bimaculatus* (42), a straightforward test for a directly comparable *osk* function in this fruit fly is not possible. However, although *D. melanogaster* mushroom body stem cells are absent in adults, analogous mushroom body neuroblasts remain mitotically active late into pupal development (43). Thus, it will be interesting to test whether *osk* functions in these neuroblasts during larval and/or pupal stages. We note that an insertion of an enhancer trap transposable element over 3 kb upstream of the *osk* transcription start site was recovered in an insertional mutagenesis screen for long term memory in in *D. melanogaster* (8). However, this insertion has not been confirmed as compromising the sequence or function of the *osk* locus, nor has *osk* been tested directly to confirm a potential role in *D. melanogaster* learning or memory.

Although *D. melanogaster* lacks adult mushroom body neuroblasts, it is possible that *osk* could function in fruit fly olfactory long-term memory in a neuroblast-independent manner. A recent study of the mushroom body output neurons has suggested that long-term memory involves the activity-dependent de-repression of mRNAs localized to granules containing Pumilio, Staufen, and Orb (oo18 RNA-binding protein) proteins (9). Given that Osk nucleates similar granules containing these proteins in the *Drosophila* oocyte (44, 45), and Osk’s ability to nucleate phase-transitioned granules in *D. melanogaster* cells(46), it would be interesting to test whether Osk is involved in the formation and/or activity of these granules in the brain. Given our recent observations of highly similar molecular interactions of conserved molecules and animal germ lines and neural cells (14), future studies could test whether additional, traditionally known “germ line” genes other than *vasa* and *piwi*, including for example *staufen* (47) and *tudor* (48), also function in *G. bimaculatus* adult neuroblasts, which would suggest that *osk* acts with conserved molecular partners in different cellular contexts. Because mushroom bodies derived from neuroblasts are a conserved arthropod brain structure across and beyond insects (49–52), it seems unlikely that *oskar* played a role in the evolution of insect mushroom body neurogenesis *per se*.

Both germ cells and neuroblasts are stem cells that give rise to highly specialized daughter cells, while remaining proliferative for long periods of time. Thus, the original role of *osk* in both cell types could conceivably be related to stem cell maintenance and/or asymmetric division. Indeed, many different highly conserved “germ line genes” including *vasa*, *nanos*, and *piwi* are found in a variety of multipotent cells in diverse animals(14, 53), raising the possibility that such genes were involved in establishing multipotency rather than specifying germ cell fate *per se*. In our previous examination of the distribution of *osk* orthologs across insects, we observed that crickets are not the only insects reported to express *osk* in the brain, as *osk* transcripts are detected in transcriptomes from the brains of cockroach, wasp and beetle species as well (16). This is consistent with the hypothesis that *osk* played an ancient neural role of some kind in insects. We previously showed evidence supporting the hypothesis that evolved changes in the biophysical characteristics of Oskar protein may have driven the evolution of a novel mechanism of germ line specification in the holometabolous insects (16). A broader understanding of the putative ancestral and derived function(s) of *osk* thus requires additional studies of phylogenetically diverse insects, as well as further detailed biochemical analysis in the context of *Drosophila* germ cells and neurons.

Our results provide an example of how newly arisen genes may find stable homes in pre-existing genetic regulatory circuits. In the case of *osk*, we hypothesize that by evolving or acquiring binding sites responsive to the conserved transcription factor Creb, *osk* may have gained expression in the brain, opening the door for potential participation in neural roles. An alternative hypothesis to *de novo* evolution of CRE sites to explain *osk*’s expression in the cricket brain, is that *osk* inherited CRE binding sites from an ancestral sequence that contributed to *osk*’s genesis. Our previous work suggested that *osk* arose through a fusion of a eukaryotic LOTUS domain in the 5’ position, coding for *osk*’s N-terminal LOTUS domain, and a prokaryotic SGNH hydrolase-like domain in the 3’ position, coding for *osk*’s C-terminal OSK domain (54). In this scenario, a pre-existing LOTUS-domaining gene could have donated not just its LOTUS domain, but also some upstream 5’ regulatory sequences, including CRE sites and/or neural expression elements, to *osk*. This hypothesis might predict that extant LOTUS domain-containing genes might display one or both of CRE binding sites, or expression in the brain. To test this hypothesis, we searched the *G. bimaculatus* genome for LOTUS domain-containing genes, and identified five such genes (Supplementary File 2). These were *osk*, *Tdrd5*, *Tdrd7*, *limkain b1*, and an uncharacterized gene with annotation ID GBI_15344 (Supplementary File 2). In transcriptomes previously generated from the adult brain (55), we detected levels of all four non-*oskar* genes at levels at least as high as that detected for *osk* (Suppl. Figure S4, Suppl. Table S8). Moreover, in the 10kb upstream of the first predicted codon of these genes, we detected a putative CRE binding site (Suppl. Table S9). With the caveat that our transcriptomes do not provide spatial or cell-type resolution for the expression data, both of these findings are consistent with the hypothesis that whatever eukaryotic LOTUS domain-containing sequence was the ancestor of *osk*’s LOTUS domain, also contributed one or both of CRE-responsive or brain-expressed upstream regulatory sequences to *osk*. Future studies will be needed to elucidate the molecular mechanisms of *osk* gene products in the cricket brain, and specifically in learning and memory.

We further speculate that the biophysical properties of Osk protein that make it effective at sequestering RNAs and participating in translational control in the germ line(56–58), may have been advantageous in promoting the rapid translation needed the synaptic plasticity that underlies learning and memory. These include Osk’s ability to form phase transitioned condensates (46, 59), its regions of high predicted disorder(16, 46, 59), and its ability to achieve and maintain asymmetric subcellular localization, all of which are well known in the germ line, and may have provided a selective advantage to Osk in the context of promoting neuronal function.

## Materials and Methods

*Gryllus bimaculatus* husbandry, in situ hybridization, immunostaining, olfactory learning assays, RNAi and qPCR were performed as previously described. See Extended Materials & Methods in Supporting Information for references and detailed protocols.

### EdU Assay

Cell proliferation was assayed using the Click-iT EdU Alexa 488 kit (Life Technologies, Cat# C10637). Crickets were injected with 10-15μl of EdU either into the abdomen or into the head capsule through the median ocellus (both methods successfully labeled dividing neuroblasts), and brains were dissected to visualize EdU incorporation four hours post-injection. Brains were dissected and de-sheathed in ice-cold 1x PBS. Calyces were removed with a microscalpel and incubated in 0.1M citric acid for 15-30 minutes on a poly-lysinated slide (Sigma Aldrich, Cat. No. P8920-100ML). Calyces were then spread into a monolayer by adding a Sigma cote-covered coverslip, and the entire slide was flash-frozen in liquid nitrogen. The coverslip was removed, leaving the mushroom body monolayer on the slide. Slides were air-dried and were then fixed for 15 minutes in 4% PFA. EdU detection was then carried out following manufacturer’s instructions. EdU-positive cells were photographed under epifluorescence on a Zeiss AxioImager Z.1 compound microscope using Zen, and manually quantified in ImageJ. For any mushroom body where the EdU-positive cluster of cells was damaged or destroyed during preparation, that sample was discarded and not included in the analysis. For tissue double stained to visualize transcripts and EdU incorporation simultaneously, the in situ hybridization was conducted before the visualization of incorporated EdU. AxioImager Z.1, LSM 780 or LSM 880 confocal microscopes (Zeiss) were used for microscopy, driven by AxioVision or Zen (Zeiss).

### *Construction, sequencing and analysis of small RNA libraries from* G. bimaculatus *adult brains*

Unmated adult male crickets within one week of their final molt to adulthood were injected with dsRNA as described above (see *RNA interference*). At 48h post RNAi injections, brains (Suppl. Table S2) were dissected in ice-cold 1x PBS, and transferred into Trizol, following which total RNA was extracted from them following manufacturers protocols. Next, RNA was size selected for 18-30nt size range after denaturing PAGE. A 2S rRNA specific oligo was used for 2S rRNA depletion. The small RNAs were ligated at the 3’ and 5’ ends by the respective adapters and purified by denaturing PAGE after each ligation. PCR was performed after reverse transcription. The PCR product was gel purified from an agarose gel to obtain the final library. The libraries were sequenced using Illumina NextSeq500 1×75bp. The resulting data (Suppl. Table S3) were uploaded onto NCBI SRA database and are publicly available under the BioProject ID PRJNA837371.

### *Identification of* G. bimaculatus *Creb genes and construction of Creb phylogenetic tree*

Putative orthologs of *Creb/ATF* family members from several animal species were initially identified by BLAST searches (Suppl. Table S5) and then downloaded from NCBI. These sequences were then used to search for putative *G. bimaculatus CrebA* orthologs in the G*. bimaculatus* genome(36). All identified sequences were then aligned with MAFFT (v 7.510)(60). A maximum likelihood tree was created in RAxML using the PROTGAMMAWAG model (61), and plotted with the FigTree package v1.4.4 (http://tree.bio.ed.ac.uk/software/figtree) (Fig.3B).

### *Bioinformatic prediction of CRE sites in* osk *and* piwi *upstream regulatory regions*

A position frequency matrix (PFM) for the full <u>c</u>yclic AMP <u>r</u>esponse <u>e</u>lement (CRE) octameric palindrome was obtained from the JASPAR database (an open source database for transcription factor binding sites(62) (Suppl. File 1). In addition to CRE, PFMs for the TATA box were also obtained from the same database. We included TATA box proximity among our search criteria for putative CRE sites, since TATA boxes are often a feature of functional promoters, and functional promoter-proximal CRE sites are reported as often occupied by Creb. These raw PFM data (Suppl. File 1) were then used as an input in FIMO (<u>F</u>ind <u>I</u>ndividual <u>M</u>otif <u>O</u>ccurrence) in the MEME suite (a motif-based sequence analysis tool (63)), and up to 10Kb of the gene sequences upstream of the predicted transcription start site for each of *G. bimaculatus oskar*, *piwi*, and *vasa* was scanned for the presence of the CRE and TATA motifs using the annotated *G. bimaculatus* genome (36), and using p<0.0001 as the stringency criteria. For comparison, *G. bimaculatus beta actin, alpha tubulin* and *FGFR* loci were subjected to the same analyses (also with p<0.0001 as stringency criteria) to assess the possibility that any randomly chosen *G. bimaculatus* gene would be predicted to have CRE sites in the 10Kb region upstream of their transcription start site using this method (Suppl. File 2). We found that for the latter three genes, there were no CRE predictions in their upstream regions (up to 10Kb from the transcription start site). Further, we bioinformatically generated one thousand 10Kb long DNA fragments of random sequence using the “random DNA sequence” tool in the Sequence Manipulation Suite (64) and then tested them for CRE prediction. Our results indicate that a CRE site is expected to occur in a randomly generated sequence at a frequency of ~1.8 CRE sites for every 10Kb tested (Suppl. Files 1,2).

### *Bioinformatic analysis of LOTUS domain-containing genes in the* G. bimaculatus *genome*

We searched the annotated *G. bimaculatus* genome (36) for genes whose protein product was predicted to contain PFAM motif PF12872, corresponding to the LOTUS domain. This search retrieved five genes (GBI_01840 “*oskar*”, GBI_13502 “*TDRD5/tejas*”, GBI_15344 “*uncharacterized*”, GBI_15604 “*limkain b1*”, GBI_03370 “*TDRD7/tapas*”). We assessed expression of these genes in brains and gonads using previously published RNA-seq libraries (55) available at NCBI (PRJNA564136). We analyzed the RNA-seq data as in (16), including removing adapters and reads shorter than 20 nucleotides with Cutadapt v3.4 (65) and quantifying the gene expression in transcripts per million (TPM) with RSEM v1.2.29 (66), using STAR v2.7.0e1 (67) as read mapper against the *G. bimaculatus* genome (36) (Suppl. Table S8). For each of these genes we retrieved and searched the 10Kb upstream of the first codon annotated for CRE sites as described above in “*Bioinformatic prediction of CRE sites in* osk *and* piwi *upstream regulatory region”* (Suppl. Table S9).

### PCR amplification, sequence confirmation and cloning of CRE sites

Based on bioinformatic predictions of putative CRE sites, primers were designed in the upstream regulatory regions of *osk* (Suppl. Table S7 #1 and #2). Once both CRE sites were sequence confirmed by Sanger sequencing, the ~30bp fragments containing each CRE site were synthetically generated as duplexes (with 3’A overhangs) for use as EMSA pre-probes (Suppl. Table S7; CRE site in bold). The 3’A overhangs were then used to clone all EMSA pre-probes into a pGEM-T easy vector following manufacturer’s instructions (Promega, catalog number A1360) using One-Shot chemically competent TOP10 *E. coli* cells (Thermo-Fisher, catalog number C4040-06).

### Generation of 5’Cy5 labelled Electrophoretic Mobility Shift Assay (EMSA) Probes and EMSA

Once cloned, pGEM-T easy specific duplex forward primer (5’Cy5-ACGTCGCATGCTCCCGGCCATG, reverse complement 5’Cy5-CATGGCCGGGAGCATGCGACGT), and reverse primer (5’Cy5-GTCGACCTGCAGGCGGCCGCGAATT, reverse complement 5-Cy5-AATTCGCGGCCGCCTGCAGGTCGAC) were designed with 5’Cy5 modifications to amplify inserts and generate fluorescently labeled double stranded EMSA probes, using a two-step PCR program with the following conditions: (98°C for 60 seconds (x1cycle); 98°C for 15 seconds followed by 72°C for 30 seconds (x30 cycles); 72°C for 5 minutes (x1cycle) (Suppl. Table S7). The PCR product was loaded onto a 1% agarose gel and the desired bands were gel eluted following IBI Scientific’s PCR purification and gel elution kit (catalog number IB47030) in 30μl water. A second round of PCR amplification following the conditions described above was performed using the eluted DNA from previous steps to increase probe yield. All steps starting with the first round of PCR were done in the dark to protect fluorescently labeled probes. Probe concentrations were measured using Nanodrop, and diluted to a final concentration of 40 fmol/probe for use in EMSA(68). 20% native PAGE gels were used to study gel shifts. Gels were imaged using an Azure Sapphire Biomolecular Imager (VWR).

### *Nuclear Protein extracts from unmated adult male* G. bimaculatus *brains*

Brains were dissected from unmated *G. bimaculatus* males within one week of their final molt that were anesthetized briefly on ice prior to dissection in 1x PBS. Nuclear protein extracts were prepared from dissected brains following manufacturer’s instructions (Abcam Nuclear Extraction Kit, catalog number ab113474).

## Acknowledgments and Funding Sources

Thanks to Taro Mito (University of Tokushima, Japan) for advice on cricket brain dissection; Venkatesh Murthy (Harvard University) for allowing us to use his laboratory’s vibratome; Elena Kramer and Min Ya (Harvard University) for help with polyacrylamide gel electrophoresis setup for EMSAs; William E. Theurkauf (University of Massachusetts Chan Medical School) for guidance on cricket small RNA library preparation and sequencing, and for financial support of SSP; members of the Extavour lab for discussion; and the Faculty of Arts and Sciences Bauer Molecular Biology Core Facility at Harvard University for Illumina Sequencing. This study was supported by National Science Foundation award #IOS-0817678, funds from Harvard University and from the Howard Hughes Medical Institute to CGE, a National Science Foundation Graduate Research Training Fellowship to BEC, and a Grant-in-Aid for Scientific Research from the Ministry of Education, Science, Culture, Sports, and Technology of Japan (no. 21K19245) to MM.

