## Supplementary Figures S1-S5, Table S1-S9, Files S1-S2 for "*oskar* acts with the transcription factor Creb to regulate long-term memory in crickets"

**This PDF file includes:**

Supporting text: Extended Materials & Methods
Figures S1 to S5
Tables S1 to S9
Supplementary File 1
Download Link and Legend for Supplementary File 2
SI References

**Other supporting materials for this manuscript include the following:**

Supplementary File 2

**Extended Materials & Methods**

**G. bimaculatus *husbandry***
For behavior experiments, *G. bimaculatus* crickets were maintained in the Mizunami laboratory at 27ºC on a 12:12 light cycle, with a diet of insect food pellets, as previously described(1). For gene expression analysis, quantitative PCR, and cell proliferation experiments, crickets were maintained in the Extavour laboratory at 28ºC and 35% relative humidity on a 12:12 light cycle, with a diet of grain and cat food, as previously described(2).

***In situ hybridization***
For in situ hybridization, brains were dissected and de-sheathed in ice-cold 1x Phosphate Buffered Saline (1X PBS) as previously described(3, 4). Brains were fixed one hour in 4% paraformaldehyde in 1X PBS, followed by an additional overnight fixation in the same solution at 4ºC, or for an additional 3-4 hours at room temperature. *osk* transcripts were detected using a 788 bp probe, following standard protocols(2) with the following modifications to reduce background: 20-minute Proteinase K (Thermo Fisher Scientific, Cat# EO0491) treatment followed by a 30-minute fixation in 0.8% glutaraldehyde in 1X PBS and 4% paraformaldehyde in 1X PBS. The *osk* probe was used at 1.0 ng/µl concentration and hybridized at 69-70ºC. Brains were sectioned after in situ development was completed, by embedding in 4% low-melt agarose in distilled water, and sectioning at 50-90µM using a Leica VT1000S vibratome.

***Immunostaining***
For immunostaining, primary antibodies used were as follows: rabbit anti-Gb-Vasa and anti-Gb-Piwi(5) 1:300, mouse anti-RNA polymerase II pSer 6 Mab H5 (Covance MMS-129R) 1:100, FITC-conjugated anti-alpha Tubulin (Sigma F2168) 1:100 and rabbit anti-*Drosophila* Vasa (kind gift of Paul Lasko, McGill University) 1:500 following standard procedures as previously described(2). Goat anti-rabbit secondary antibodies conjugated to Alexa 488, Alexa 555 or Alexa 568 (Invitrogen) were used at 1:500 or 1:1000. Counterstains used were Hoechst 33342 (Sigma B2261) at 0.1 to 0.05 mg/ml and FITC-conjugated phalloidin (Sigma P5282) at 1 U/ml. For antibody staining, brains were embedded in 4% low-melt agarose in distilled water, and sectioned at 50-90µM using a Leica VT1000S vibratome, prior to incubation with the primary antibody. For analysis of apoptosis and DNA damage, brains were fixed 4h after EdU injection and sectioned via vibratome, with EdU detection performed first (using Invitrogen's Click-iT protocol), followed by antibody staining and confocal analysis.

***RNA extractions, cDNA preparation and cloning of gene fragments for production of dsRNA***
Brains from unmated male adults were dissected in ice-cold 1X PBS, then immediately homogenized in TRIzol (ThermoFisher Scientific, catalog number 15596026). Total RNA was extracted following the manufacturer's instructions, including a 30-minute DNAse treatment. 1µg of total RNA was used as template for cDNA synthesis using SuperScript III (ThermoFisher Scientific, catalog number 18-080-044) with oligo-dT primers. cDNA was diluted 1:10 prior to PCR with gene specific primers, and 2 µL of template was used per 25 µL PCR reaction. PCR products were run on a 1% agarose gel and desired bands were gel eluted following IBI Scientific's PCR purification and gel elution kit (catalog number IB47030). Then, products were cloned into Zero blunt TOPO PCR cloning kit (ThermoFisher Scientific, catalog number 450245) using electro-competent *DH5α-E* cells (ThermoFisher Scientific, catalog number 11319019).

***Identification and annotation of Piwi Proteins in* G. bimaculatus**
We previously used a *G. bimaculatus* transcriptome(6) to identify two RNA fragments corresponding to two Piwi family proteins (*piwi*: JQ434103 and *piwi-2*: KC242806.1(5, 7)). All previous published analyses of *piwi* in *G. bimaculatus* were performed with "*piwi*" (JQ434103), as only this gene showed enriched expression in embryonic germ cells(5). Since the time of our initial studies on *piwi*, we assembled and annotated an updated *G. bimaculatus* genome(8). For the present study, we therefore performed new BLAST searches to clarify the status of *piwi* orthologs in this cricket (Suppl. Fig. S5). We found both previously identified fragmented *piwi* RNA sequences within the new gene annotations(8) with gene IDs GBI_17641 (containing "*piwi*" fragment JQ434103) and GBI_07509 (containing "*piwi-2*" fragment KC242806.1) respectively. We also identified two additional putative novel *piwi*-like genes annotated in the *G. bimaculatus* genome, with gene IDs GBI_09750 and GBI_09796 (8).

Using InterProscan, we confirmed that the amino acid sequences of the proteins encoded by these four putative *piwi* genes contained the typical characteristics of the Argonaut/Piwi proteins(9) namely a Paz domain followed by a C-terminal Piwi domain. Additionally, we inferred the gene tree of the Argonaute/Piwi protein family using the putative *G. bimaculatus* Piwi and Argonaute protein amino acid sequences obtained from the genome, together with sequences of publicly available Piwi and Argonaute proteins from other insects (*Drosophila melanogaster*, *Apis mellifera*, *Bombyx mori*, *Tribolium castaneum*, *Blattella germanica*, *Zootermopsis nevadensis*, *Acyrthosiphon pisum*, and *Locusta migratoria)*. Protein sequence alignments were performed with MUSCLE(10, 11) in Geneious (v3.8.425; www.geneious.com), and the gene tree was inferred with RAxML v8.2.11(12) with 100 bootstrap iterations to obtain the support values of each node. The tree was then visualized with ggtree(13, 14). The resulting tree differentiated four groups of sequences with bootstrap values above 90%, each of which contained different Piwi/Argonaute subfamilies as follows: Argonaute 1 proteins (AGO1; included GBI_02015), Argonaute 2 (AGO2; included GBI_13717), Argonaute 3 (AGO3; included GBI_01357), and Piwi proteins and their paralogs (Aub and Siwi, the Piwi paralogs in *D. melanogaster* and *B. mori* respectively; included GBI_17641) (Suppl. Fig. S5). This indicated that our previous analyses(5) had indeed targeted the true *piwi* ortholog in *G. bimaculatus*. Accordingly, all gene expression and function analyses in the present study were also performed on this true *piwi* ortholog (GBI_17641).

Analysis of the small RNA species present in $piwi^{RNAi}$ animals indicated that our *piwi* RNAi experiments specifically targeted the true *piwi* (GBI_17641), and did not impact expression of the other 3 *piwi* subfamily genes (Suppl. Table S4). The quality control of the eight small RNA-seq samples (3 $piwi^{RNAi}$, 2 $osk^{RNAi}$, 2 $DsRed^{RNAi}$, and 1 untreated or wild-type, WT) was performed with FastQC v0.11.8(15), and the adapters were trimmed with Cutadapt v1.8.1(16). Clean reads were mapped to the *G. bimaculatus* genome assembly(8) with Bowtie 2 v2.3.4.1(17) using parameters "-L 18, -N 0". The numbers of sequenced reads, clean reads, and mapped reads are shown in Suppl. Table S1. The mapped reads were retrieved using samtools v1.9(18) for obtaining the read length distributions (Suppl. Fig. S1B). The proportion of miRNAs and piRNAs in each sample was extrapolated as the percentage of reads of 22-23 nts and 28-29 nts respectively (Suppl. Fig. S1B). The FeatureCounts function from the R package Rsubread v2.0.0(19) was used to count the number of reads mapped to all annotated genes and build a table of counts. The counts of o*sk* (GBI_0140) and the four *piwi* genes (GBI_09750, GBI_09796, GBI_07509, and GBI_17641) were retrieved (Suppl. Table S4). The small RNA-seq reads mapping to the target genes were assumed to be reads of the siRNAs produced from the dsRNA(20). The thousands of such reads mapping to our targeted *piwi* (GBI_17641) and the absence of such siRNA reads mapping to other p*iwi* genes, suggest that no off-target effects impaired the expression of other *piwi* genes.

***RNA interference (RNAi)***
Unmated adult male crickets within one week of their final molt were injected with 2 µL of double-stranded RNA (dsRNA) through a hole pierced in the median ocellus(21) using a 10 µL syringe fitted with a 26S gauge tip (WPI, Tokyo, Japan; Hamilton Inc., Nevada, USA). Behavioral tests were repeated using two non-overlapping fragments of *oskar* (742bp and 503bp), a 646bp fragment of *piw*i, a 541bp fragment of *vasa*, two non-overlapping fragments of *CrebA* (both 387bp fragments), a 384bp fragment of *CrebB*, and a 678bp fragment of *DsRed* as a negative control (7). Double-stranded RNA concentrations used were 10 µM for *oskar*; 3.38 µg/µL for *piwi*; 2.71 µg/µL for *vasa*; 2.97µg/µL for *DsRed*; 6 µg/µL for *oskar/piwi* double knock down*;* 7µg/µL for *CrebA;* 7µg/µL for *CrebB; and* 7µg/µL for *DsRed* (Suppl. Table S2).

***Gene expression from mRNA-seq data***
To check the expression of *G. bimaculatus* genes in nervous systems in wild-type animals, previously generated mRNA-seq libraries were used(22) and the complete CDS for genes of interest were obtained from the recently published genome[33]. Reads were trimmed with Cutadapt v3.4(16), and mapped to the full *G. bimaculatus* CDS using Geneious Read Mapper(23). DESeq2 normalized counts in fragments per kilobase per exon per million mapped fragments (FPKMs) were then obtained for all genes of interest. The expression of *osk* and *piwi* genes (Suppl. Table S4) shows that the depleted *piwi* (GBI_17641) is one of the two *piwi* genes expressed in wild-type brains of males and females. In the same way, we obtained the expression in FPKMs of *vasa* (GBI_17344), *CrebA* (GBI_04244), and *CrebB* (GBI_02305) (Suppl. Table S6).

***Olfactory learning behavioral memory assays***

Adult male crickets at eight days after the final molt were used in all experiments, because learning and memory capabilities are highly affected by reproductive status and aging (24), and in our experience, young unmated males exhibit more stable memory scores and longer memory retention scores than females or older males (unpublished observations). Three days before conditioning, individual crickets were separated into 100-mL beakers and deprived of drinking water to enhance their motivation to search for water. Two days before conditioning (ten days after the imaginal molt), each cricket was injected with dsRNA as described above. Two days after dsRNA injection, each cricket was subjected to an odor preference test, in which the animal was allowed to freely visit peppermint and vanilla odors(25). The time spent at each of the peppermint and vanilla odor sources was measured cumulatively to evaluate relative odor preference(25). Crickets were subjected to 4-trial conditioning, in which an odor was paired with water reward, with an inter-trial interval of five minutes(4, 26). For conditioning, a small filter paper was attached to the needle of a hypodermic syringe. The syringe was filled with water reward (unconditioned stimulus), and the filter paper was soaked with peppermint essence (conditioned stimulus). At one hour and one day after the end of the conditioning, each cricket was subjected to an odor preference test. The relative odor preference of each conditioned and control animal was measured using the preference index (PI) for rewarded odor (peppermint), defined as tP/(tP+tV) *100 (%), where tP is the time spent exploring the peppermint source and tV is the time spent exploring the vanilla (unrewarded) odor. Wilcoxon's test was used to compare odor preferences before and after training. For multiple comparisons, Holm's method was used to adjust the significance level.

***Quantitative PCR (qPCR)***

Two days after dsRNA injection, brains were dissected from unmated male adults (within a week of final molt to adulthood) in ice-cold 1x PBS, then immediately homogenized in TRIzol (ThermoFisher Scientific, catalog number 15596026). Total RNA was extracted from a total of six brains per treatment, following the manufacturer's instructions, including a 30-minute DNAse treatment. 1 µg of total RNA was used as template for cDNA synthesis using SuperScript III (ThermoFisher Scientific, catalog number 18-080-044) with oligo-dT primers. cDNA was diluted 1:10 prior to qPCR, and 6 µL of template was used per 25 µL qPCR reaction. (PerfeCta SYBR Green SuperMix, Low ROX, Quanta Biosciences, catalog number 101414-158). qPCR reactions were conducted in triplicate, and fold change was calculated using the $\Delta\Delta$Ct method(27), with standard deviation propagated following standard methods. *Beta-tubulin* was used as a reference gene(2). Primers amplifying a 130bp fragment of *Gb-CrebA*, a 140bp fragment of *Gb-CrebB*, a 234bp fragment of *Gb-oskar*, a 166 bp fragment of *Beta-tubulin* (2), a 129bp fragment of *Gb-piwi*, a 150bp fragment of *Gb-vasa* and a 120bp fragment of *Gb-FGFR* (Fibroblast Growth Factor Receptor) were used (Suppl. Table S1).

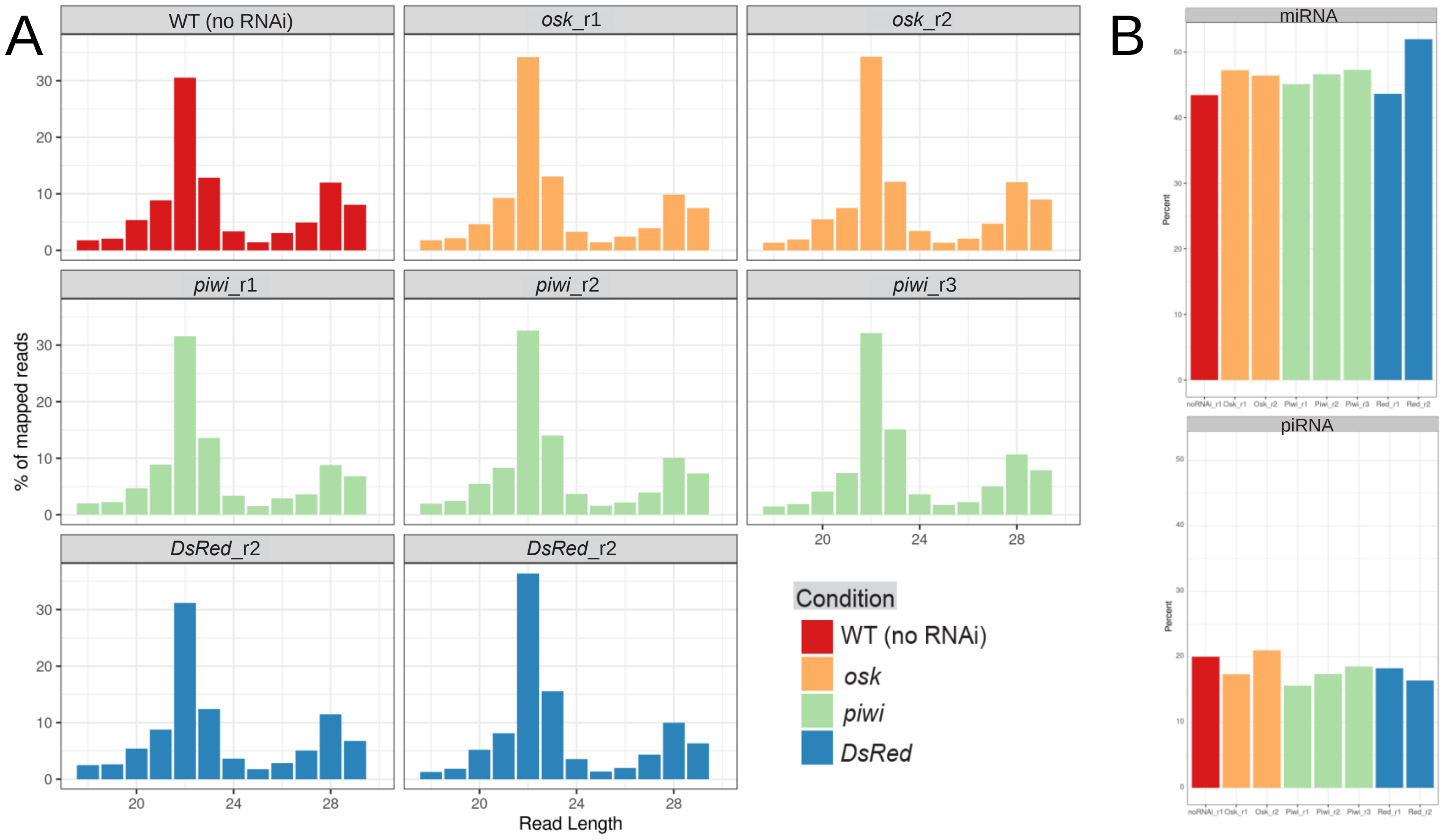


**Figure S1. Small RNA library mapping and analysis (A)** Read Length Distribution**:** Percentage of mapped reads of each length from 18 to 29 nucleotides in each sequenced small RNA library colored by RNAi treatment. The two peaks at ~22 and ~28 mainly correspond to miRNAs and piRNAs. **(B)** miRNAs vs piRNA: Estimated proportion of reads corresponding to microRNAs and piRNAs in each sample (colored by condition) based on the percentage of reads of 22-23 nucleotides in length and 28-29 nucleotides respectively.

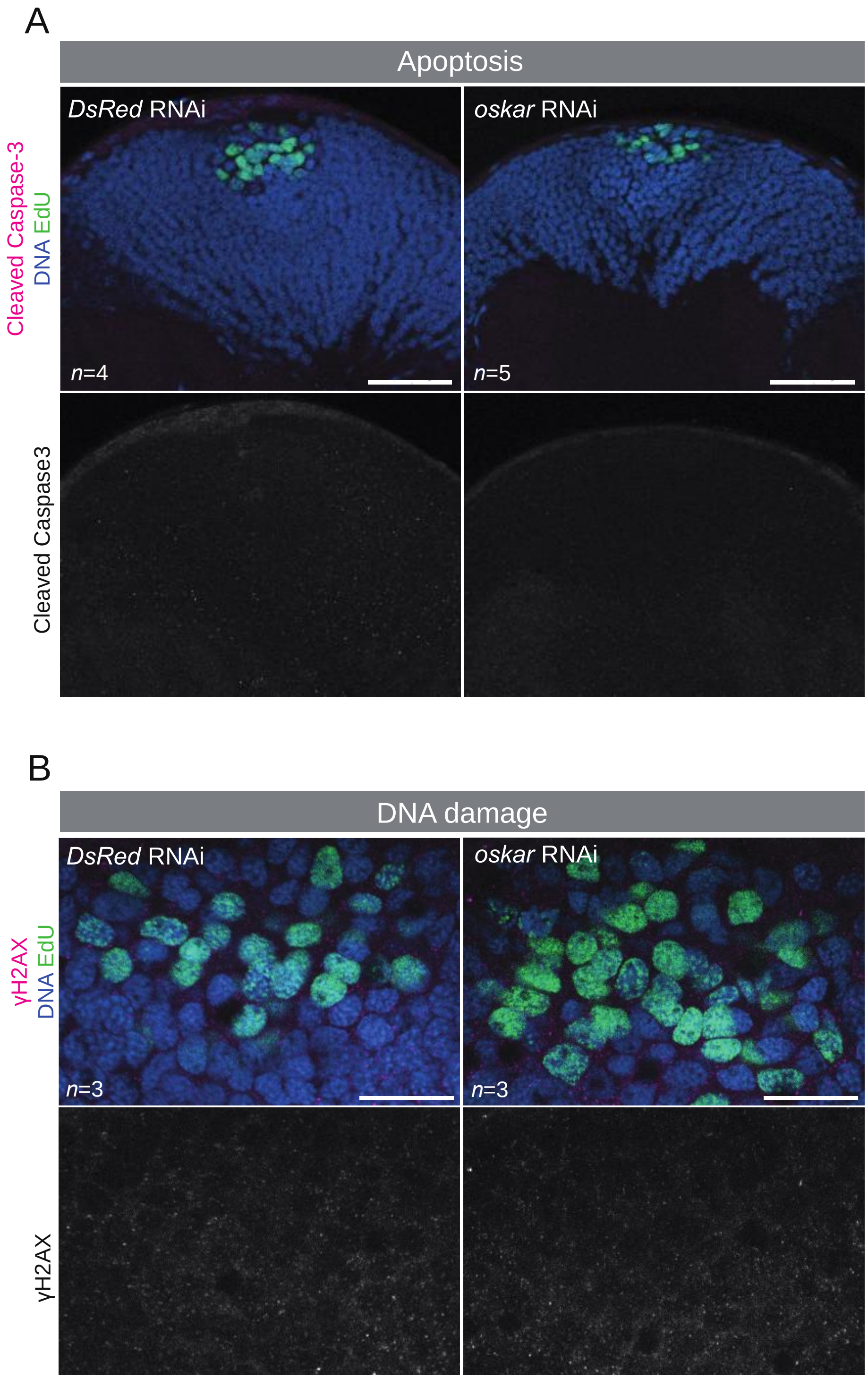


**Figure S2. Assessment of DNA damage and apoptosis in *osk*$^{RNAi}$ adult mushroom body neuroblasts.** Apoptosis marker Cleaved caspase 3 (**A**) and DNA damage marker gamma H2AX immunostaining (**B**) in adult mushroom bodies, including neuroblasts of control and *osk*$^{RNAi}$ brains.

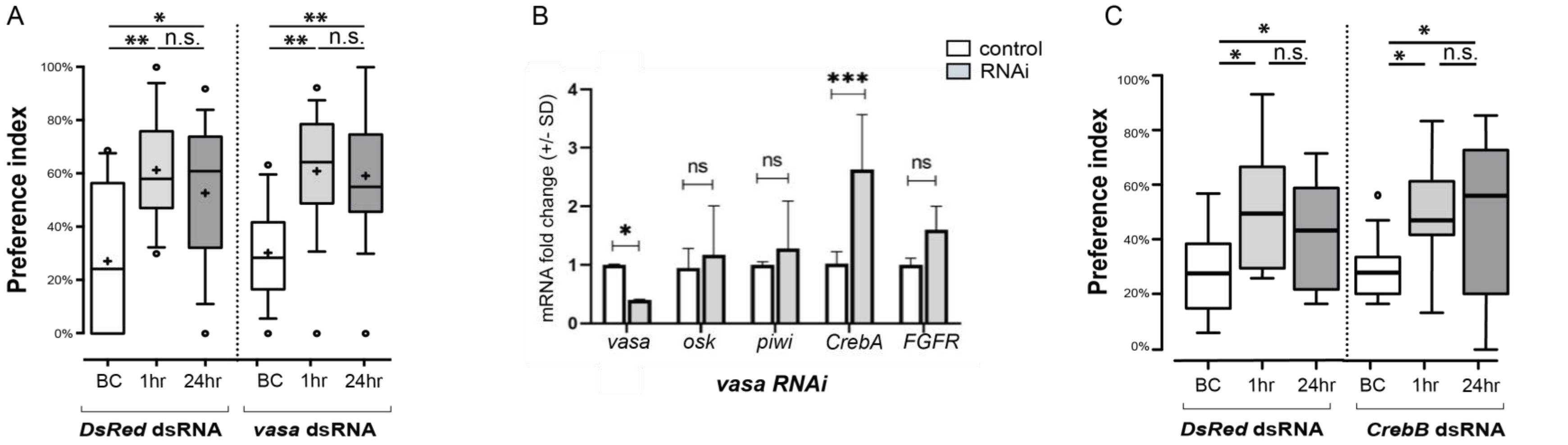


**Figure S3. Vasa and CrebB are not required for long-term memory in the cricket *G. bimaculatus*.** (**A**) *vasa* RNAi fails to recapitulate the long-term olfactory memory phenotype seen in *osk* and *piwi* RNAi. BC = "Before Conditioning", 1hr = "1 hour post training" and 24hrs = "24 hours after training". (**B**) qPCR on $vasa^{RNAi}$ brains shows significant up-regulation of *CrebA*. (**C**) $CrebB^{RNAi}$ does not recapitulate the long-term memory phenotype shown by $CrebA^{RNAi}$*.* N=9 for *CrebB,* and N=10 for *DsRed.* (* p < 0.05, ** p < 0.01, *** p<0.001, **** p < 0.0001, n.s. = not statistically significant).

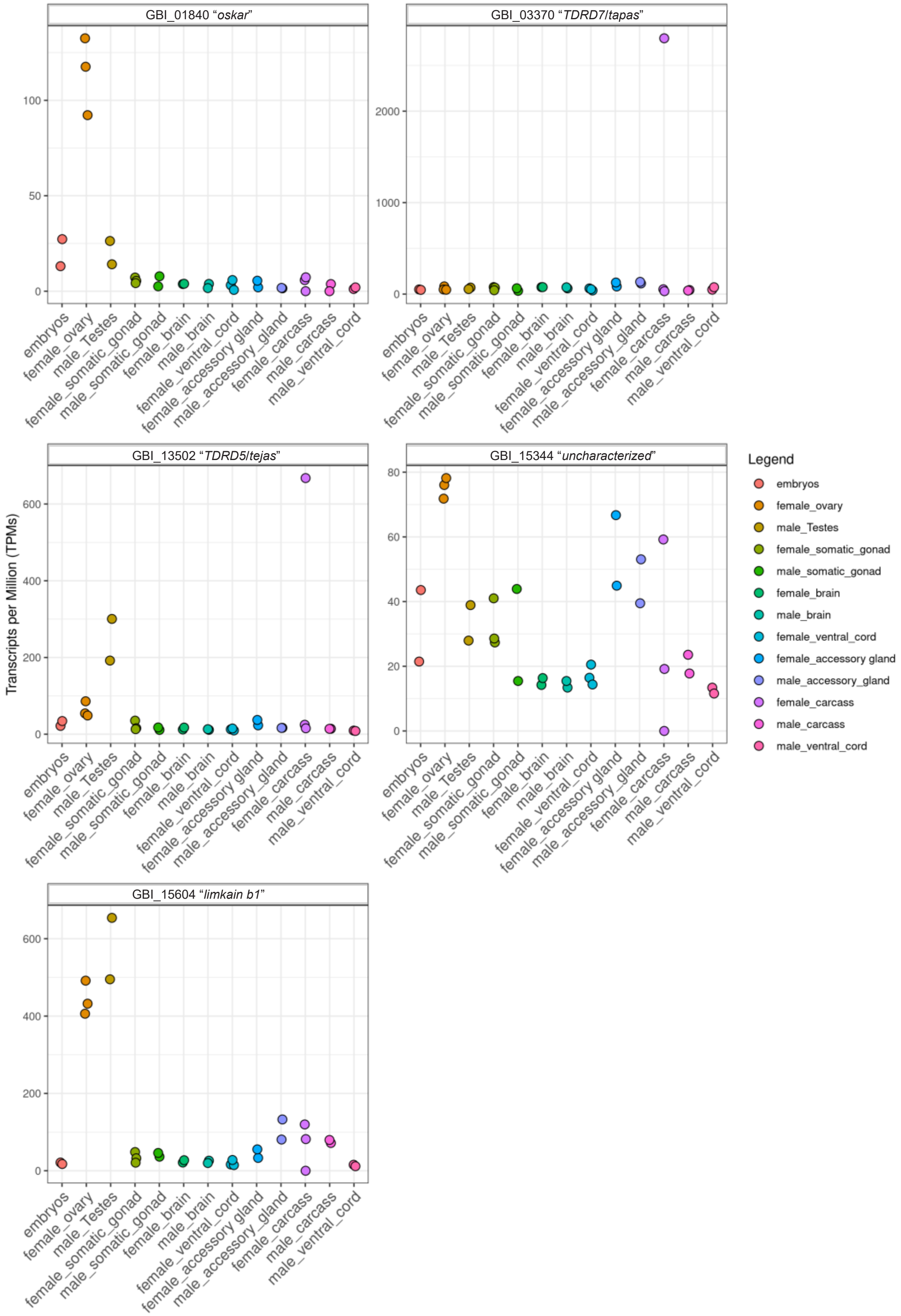


**Figure S4. LOTUS domain-containing gene expression in *G. bimaculatus***. Expression plots (in transcripts per million (TPM)) for LOTUS-domain containing genes in *G. bimaculatus* brain and gonad transcriptomes (corresponds to data shown in Table S8).

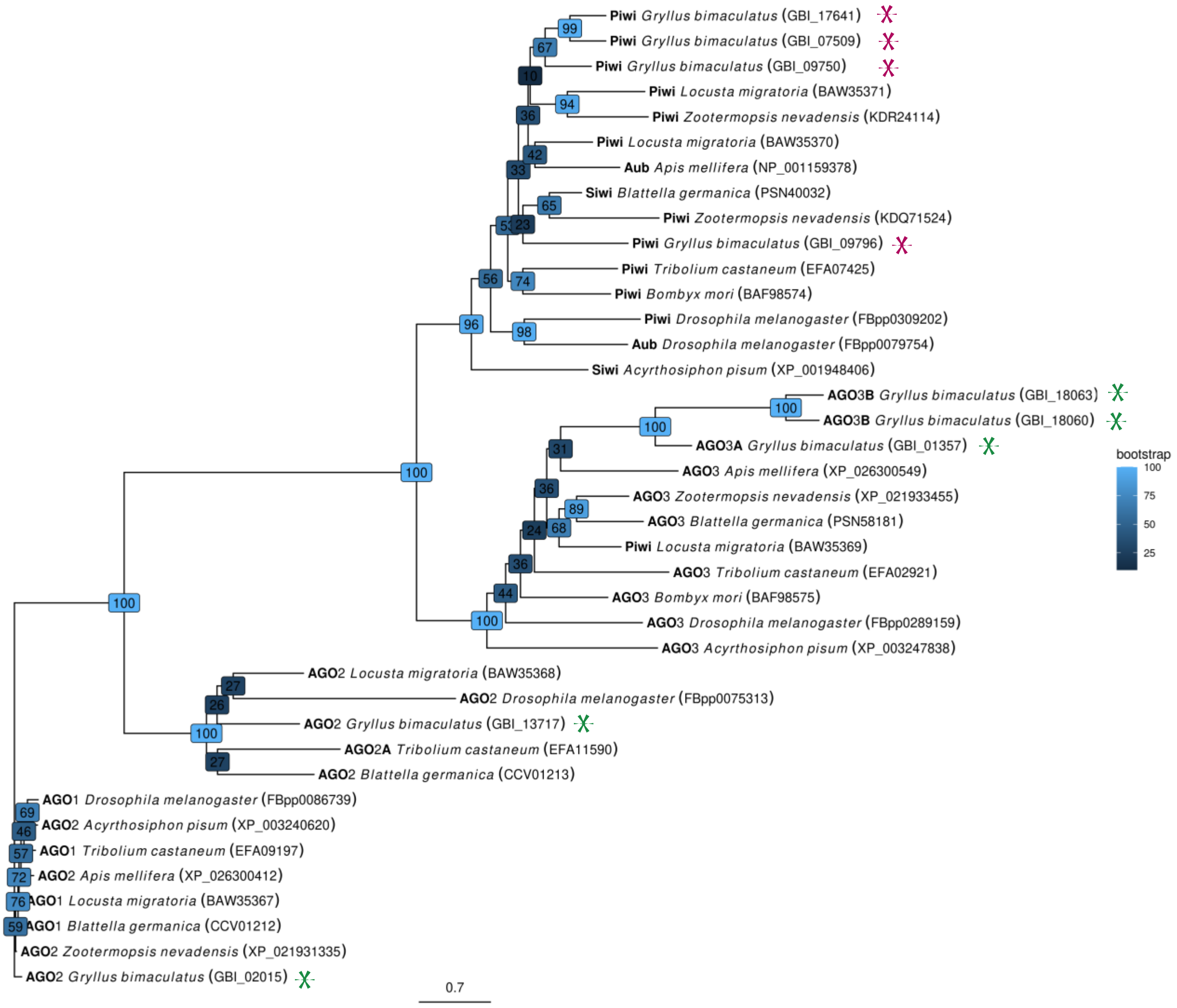


**Figure S5. Piwi family genes in the cricket *G. bimaculatus*.** *G. bimaculatus piwi* ortholog identification and phylogenetic analysis. Argonaute family gene tree generated with the PIWI, AUB, AGO1, AGO2, and AGO3 protein sequences from *Drosophila melanogaster*, *Apis mellifera*, *Bombyx mori*, *Tribolium castaneum*, *Blattella germanica*, *Zootermopsis nevadensis*, *Acyrthosiphon pisum*, and *Locusta migratoria*. Values at nodes represent bootstrap support, in boxes color-coded from dark (lowest) to light (highest) blue. *G. bimaculatus piwi* and *argonaut* genes indicated by red and green asterisks respectively.

| Gene | qPCR Primers |
|---|---|
| *CrebA* | F:CCGCCTTCACCACCGCAGAC |
| | R:ATGCTTAGTTTGGGGATGACGACGC |
| *CrebB* | F:AGACTCCTGCTAATATTCAGCCTGT |
| | R:TAGACTGTCATCACTTCCTGCTTCT |
| *piwi* | F:TTCGGCCAACTACTTCAAGC |
| | R:AGAGTTTCCCGATGAACACG |
| *vasa* | F:GAACATTGTGAGCCTCATGC |
| | R:TTGCTGAGCCTGGTGGTAT |
| *oskar* | F:TTGTTGACCATTCCCTTCCT |
| | R:ACTCCACAACACCACTCC |
| *Beta-Tubulin* | F:TGGACTCCGTCCGGTCAGGC |
| | R:TCGCAGCTCTCGGCCTCCTT |
| *FGFR* | F:ACCTGTCTTCAGCGAACTAGTG |
| | R:ACTTGCTTCTTGGCTGGATG |

**Table S1**. Primers used for quantitative PCR of all listed *G. bimaculatus* genes. All sequences are in 5’ to 3’ orientation.

| Biological replicate # | RNAi Experimental condition | # adult males injected | µl dsRNA injected | dsRNA concentration |
|---|---|---|---|---|
| 1 | Control 1 (uninjected) | 16 | N/A | N/A |
| | Control 2 (*DsRed* injected) | 13 | 4 | 4.4µg/µl |
| | *piwi* | 17 | 4 | 4.6µg/µl |
| 2 | Control 2 (*DsRed* injected) | 10 | 3 | 10 µg/µl |
| | *piwi* | 17 | 3-4 | 8 µg/µl |
| | *oskar* | 14 | 4 | 7.8-14µg/µl |
| 3 | Control 2 (*DsRed* injected) | 14 | 3 | 9.8µg/µl |
| | *piwi* | 11 | 3 | 10µg/µl |
| | *oskar* | 16 | 3 | 11µg/µl |

**Table S2**. A total of 128 unmated, adult male cricket brains (16 brains from Control 1 "uninjected controls", 37 brains from Control 2 "*DsRed* injected", 45 brains from "*piwi*" dsRNA injected, and 30 brains from "*osk*" dsRNA injected) were dissected 48h post dsRNA injection and processed for making small RNA libraries.

| File Name | Sample | # Raw reads | # Clean reads | % Clean | # Mapped reads | % Mapped |
|---|---|---|---|---|---|---|
| Bill.smRNA.cricket_dsRed_RNAi.brain.r1.fastq.gz | DsRed_RNAi.brain.r1 | 9,193,785 | 4,404,190 | 47.90% | 3,568,586 | 81.03% |
| Bill.smRNA.cricket_dsRed_RNAi.brain.r2.fastq.gz | DsRed_RNAi.brain.r2 | 36,880,382 | 30,410,536 | 82.46% | 25,931,302 | 85.27% |
| Bill.smRNA.cricket_No_RNAi.brain.r1.fastq.gz | No_RNAi.brain.r1 | 11,311,618 | 6,373,759 | 56.35% | 5,317,264 | 83.42% |
| Bill.smRNA.cricket_osk_RNAi.brain.r1.fastq.gz | osk_RNAi.brain.r1 | 10,811,667 | 6,439,713 | 59.56% | 5,086,605 | 78.99% |
| Bill.smRNA.cricket_osk_RNAi.brain.r2.fastq.gz | osk_RNAi.brain.r2 | 29,321,677 | 24,047,151 | 82.01% | 20,518,912 | 85.33% |
| Bill.smRNA.cricket_piwi_RNAi.brain.r1.fastq.gz | piwi_RNAi.brain.r1 | 7,120,996 | 3,637,311 | 51.08% | 2,828,184 | 77.75% |
| Bill.smRNA.cricket_piwi_RNAi.brain.r2.fastq.gz | piwi_RNAi.brain.r2 | 18,492,553 | 13,202,708 | 71.39% | 10,629,195 | 80.51% |
| Bill.smRNA.cricket_piwi_RNAi.brain.r3.fastq.gz | piwi_RNAi.brain.r3 | 21,923,563 | 17,222,470 | 78.56% | 13,628,698 | 79.13% |

**Table S3**. Number of raw sequenced small RNA reads, number and percentage of clean reads, and number and percentage of reads mapped to the *G. bimaculatus* genome.

| Gene ID | Gene Name | *DsRed.r1* | *DsRed.r2* | WT(no RNAi) | *osk.r1* | *osk.r2* | *piwi.r1* | *piwi.r2* | *piwi.r3* |
|---|---|---|---|---|---|---|---|---|---|
| GBI_01840 | *oskar* | 6 | 65 | 0 | 17945 | 109776 | 6 | 2065 | 1 |
| GBI_09750 | *piwi* | 1 | 4 | 0 | 0 | 2 | 0 | 4 | 0 |
| GBI_09796 | *piwi* | 0 | 2 | 0 | 0 | 0 | 1 | 2 | 0 |
| GBI_07509 | *piwi* | 1 | 0 | 0 | 0 | 0 | 0 | 1 | 0 |
| GBI_17641 | *piwi* | 156 | 51 | 5 | 6 | 332 | 6346 | 41552 | 8940 |

**Table S4**. Number of small RNA-seq reads mapped to *osk* and *piwi* genes to assess specificity of RNAi knockdowns. These small RNA-seq reads come from the siRNA detected following injections of dsRNA for these respective genes. The targeted sequence in each library is highly enriched by small RNA-seq reads. In the *piwi*$^{RNAi}$ experiments, only the targeted *piwi* (GBI_17641) shows a high number of mapped reads in the libraries generated from animals injected with the dsRNA against *piwi*, indicating that the other three *piwi* orthologs present in the *G. bimaculatus* genome were unlikely to be targeted by our approach.

| *Mus musculus* | *Drosophila melanogaster* | *Tribolium castaneum* | *Apis mellifera* | *Acyrthosiphon pisum* | *Zootermopsis nevadensis* | *Gryllus bimaculatus* |
|---|---|---|---|---|---|---|
| OASIS (NP_036087.2) | CrebA (NP_524087.3) | CREB-A (XP_966968.2) CREB-like (XP_973089.1) | CREB-A (XP_003250132.1) CREB-like (XP_001121941.2) | CREB-A (XP_001948312.1) CREB-like (XP_001949209.1) | CREB-A (KDR23733.1) CREB-like (KDR11962.1) | CREB-like (GAIZ01013153, *Gryllus firmus* TSA), CrebA (GBI_04244, *Gryllus bimaculatus*) |
| CREB1 (NP_034082.1) CREM (NP_001104320.1) ATF1 (NP_031523.3) | CrebB (NP_001097017.1) | XP_008192794 | XP_623392.3 | XP_008186705.1 | KDR23211.1 | GAIZ01012380 and GAIZ01007852 (*Gryllus firmus* TSA), CrebB (GBI_02305, *Gryllus bimaculatus*) |
| ATF2 (NP_001020264.1) | dATF2 (NP_001033973.1) | XP_974257.1 | XP_003249317.1 | - | KDR15907.1 | - |
| ATF3 (NP_031524.2) | dATF3 (NP_620473.1) | XP_008192299.1 | XP_003251072.1 | XP_003243558.1 | KDR22659.1 | - |
| ATF4 (NP_001274109.1) ATF5 (NP_109618.1) | cryptocephal (NP_524897.1) | NP_001280506.1 | XP_006562898.1 | XP_003247514.1 | KDR14457.1 | - |
| ATF6 (NP_001074773.1) | dATF6 (NP_995745.1) | XP_008201619.1 | XP_395889.5 | XP_008183931.1 | KDR16855.1 | GAIZ01013605 (*Gryllus firmus* TSA) |

**Table S5**. GenBank IDs of *Creb/ATF* family member orthologs in mouse and other insects including the cricket *G. bimaculatus*. This information was used to construct a Creb phylogenetic tree (Fig. 3B) to infer the evolutionary relationships between mammalian Creb proteins and their insect counterparts.

| *G. bimaculatus* gene name | Gene ID | (gene expression in FKPM per tissue) | | | | | | | |
|---|---|---|---|---|---|---|---|---|---|
| | | Female brain 1 | Female brain 2 | Male brain 1 | Male brain 2 | Female ventral 1 | Female ventral 2 | Male ventral 1 | Male ventral 2 |
| *oskar* | GBI_01840 | 2.76 | 3.85 | 3.28 | 1.24 | 3.24 | 0.84 | 1.38 | 1.29 |
| *vasa* | GBI_17344 | 33.65 | 31.18 | 32.84 | 24.66 | 55.2 | 28.47 | 28.54 | 22.4 |
| *CrebA* | GBI_04244 | 198.35 | 155.47 | 170.68 | 158.7 | 258.26 | 225.45 | 218.68 | 155.47 |
| *CrebB* | GBI_02305 | 46.99 | 42.01 | 47.27 | 36.85 | 49.45 | 42.64 | 27.01 | 29.12 |
| *piwi* | GBI_09750 | 0.34 | 0.24 | 0 | 0 | 0.24 | 0.07 | 0.23 | 0.4 |
| *piwi* | GBI_09796 | 21.4 | 18.25 | 36.83 | 44.19 | 16.96 | 22.43 | 35.34 | 47.58 |
| *piwi* | GBI_07509 | 0.94 | 0.26 | 0.95 | 1.28 | 3.18 | 1.14 | 0.58 | 1.67 |
| *piwi* | GBI_17641 | 64.43 | 57.71 | 54.72 | 59.32 | 61.65 | 54.38 | 55.78 | 33.71 |

**Table S6**. Gene expression levels for *osk*, *piwi*, *vasa*, and *CrebA/B* (in FPKM per tissue) from brain and ventral cord transcriptomes of male and female adult *Gryllus bimaculatus*(22). Gene IDs as per the annotated cricket genome(8).

| CRE site # | Gene | Site | Cloning primer | EMSA Probe sequence | EMSA Probe reverse complement sequence |
|---|---|---|---|---|---|
| 1 | *oskar* | CRE-I | F: ACAGCCTGAGGCGCTATCTA | CATCCAAAGAGCG**TGGCGTCA**CGTATCAGC | GCTGATACG**TGACGCCA**CGCTCTTTGGATG |
| | | | R: AGCGTCTTCTCTGGCGACTA | | |
| 2 | *oskar* | CRE-II | F: TCCTAGCGATTTTCGCTGAC | TTATT**TTACGTCA**ATGAAACATAATTAATTCG | CGAATTAATTATGTTTCAT**TGACGTAA**AATAA |
| | | | R:TCAACTTCTCCCACATTCCA | | |

**Table S7**. Primers used for cloning and generation of EMSA probes for *G. bimaculatus oskar.* All sequences are in 5’ to 3’ orientation. **Bold face type** indicates predicted CRE site in probe sequences.

| GeneID | GBI_01840 | GBI_03370 | GBI_13502 | GBI_15344 | GBI_15604 |
|---|---|---|---|---|---|
| Gene name | oskar | TDRD7 "Tapas" | TDRD5 "Tejas" | uncharacterized | limkain b1 -like |
| GB_embryos_sample1 | 13.1 | 51.12 | 21.67 | 21.46 | 21.22 |
| GB_embryos_sample2 | 27.26 | 46.48 | 34.29 | 43.57 | 17.46 |
| GB_female_accessory gland_sample1 | 2 | 83.19 | 23.1 | 44.92 | 33.16 |
| GB_female_accessory gland_sample2 | 5.49 | 126.53 | 37.19 | 66.72 | 55.36 |
| GB_female_brain_sample1 | 3.74 | 76.1 | 12.26 | 14.22 | 21.25 |
| GB_female_brain_sample2 | 3.89 | 76.39 | 17.28 | 16.35 | 27.26 |
| GB_female_carcass_sample1 | 5.82 | 52.8 | 24.59 | 59.2 | 119.77 |
| GB_female_carcass_sample2 | 7.27 | 31.89 | 15.42 | 19.17 | 81.82 |
| GB_female_carcass_sample3 | 0 | 2796.68 | 667.6 | 0 | 0 |
| GB_female_ovary_sample1 | 132.49 | 50.54 | 54.23 | 71.82 | 406.07 |
| GB_female_ovary_sample2 | 92.23 | 47.76 | 48.75 | 78.18 | 432.07 |
| GB_female_ovary_sample3 | 117.59 | 84.3 | 85.82 | 76.04 | 491.6 |
| GB_female_somatic_gonad_sample1 | 5.41 | 70.59 | 15.6 | 27.36 | 32.25 |
| GB_female_somatic_gonad_sample2 | 4.23 | 42.27 | 13.09 | 28.58 | 20.9 |
| GB_female_somatic_gonad_sample3 | 7.16 | 76.76 | 35.41 | 41.01 | 48.4 |
| GB_female_ventral_cord_sample1 | 3.16 | 62.18 | 12.76 | 16.45 | 16.23 |
| GB_female_ventral_cord_sample2 | 0.68 | 39.68 | 9.85 | 14.37 | 14.63 |
| GB_female_ventral_cord_sample3 | 5.83 | 53.59 | 14.91 | 20.52 | 27.84 |
| GB_male_accessory_gland_sample1 | 1.49 | 116.51 | 16.58 | 53.06 | 132.66 |
| GB_male_accessory_gland_sample2 | 1.73 | 133.82 | 16.34 | 39.48 | 80.7 |
| GB_male_brain_sample1 | 3.82 | 62.6 | 11.38 | 13.41 | 26.05 |
| GB_male_brain_sample2 | 1.63 | 74.48 | 13.14 | 15.45 | 19.96 |
| GB_male_carcass_sample1 | 3.74 | 44.63 | 14.16 | 17.77 | 71.82 |
| GB_male_carcass_sample2 | 0 | 38.8 | 14 | 23.56 | 79.59 |
| GB_male_somatic_gonad_sample1 | 2.55 | 64 | 17.6 | 43.9 | 46.03 |
| GB_male_somatic_gonad_sample2 | 7.81 | 35.91 | 11.19 | 15.45 | 36.34 |
| GB_male_Testes_sample1 | 26.31 | 54.91 | 191.92 | 27.94 | 495.09 |
| GB_male_Testes_sample2 | 14.1 | 71.36 | 300.5 | 38.91 | 653.98 |
| GB_male_ventral_cord_sample1 | 1.15 | 48.55 | 9.37 | 13.39 | 15.5 |
| GB_male_ventral_cord_sample2 | 1.98 | 75.09 | 8.43 | 11.58 | 11.73 |

**Table S8.** Expression levels of LOTUS domain-containing genes (in transcripts per million (TPM)) in the *G. bimaculatus* genome (data plotted in Supplementary Figure S4).

| Sequence name | # putative CRE motifs found | putative CRE motif sequence (5' to 3') | Strand | Start | End | p-value (p<0.0001) | q-value |
|---|---|---|---|---|---|---|---|
| **TDRD5 "tejas"_GBI_13502_upstream10Kb** | 1 | TGACGYMA | plus | 3925 | 3932 | 5.93E-05 | 1 |
| **TDRD7 "tapas"_GBI_03370_upstream10Kb** | 2 | TGACGYMA | plus | 8185 | 8192 | 1.47E-05 | 0.147 |
| | | TGACGYMA | minus | 8185 | 8192 | 1.47E-05 | 0.147 |
| **uncharacterized_GBI_15344_upstream10Kb** | 3 | TGACGYMA | minus | 5868 | 5875 | 2.67E-05 | 0.534 |
| | | TGACGYMA | plus | 5769 | 5776 | 7.40E-05 | 0.573 |
| | | TGACGYMA | minus | 7355 | 7362 | 8.60E-05 | 0.573 |
| **Limkain b1_GBI_15604_upstream10Kb** | 3 | TGACGYMA | plus | 6110 | 6117 | 1.47E-05 | 0.137 |
| | | TGACGYMA | minus | 6110 | 6117 | 1.47E-05 | 0.137 |
| | | TGACGYMA | minus | 3128 | 3135 | 8.60E-05 | 0.536 |
| **oskar_GBI_01840** | 2 | TGACGYMA | minus | 8016 | 8023 | 2.67E-05 | 0.883 |
| | | TGACGYMA | minus | 6419 | 6426 | 5.93E-05 | 0.883 |
| **piwi_GBI_17641** | 2 | TGACGYMA | minus | 2503 | 2510 | 5.93E-05 | 0.474 |
| | | TGACGYMA | minus | 5048 | 5055 | 5.93E-05 | 0.474 |

| Sequence name | Matched CRE sequence (5' to 3') | Total # putative TATA motifs in 10Kb | # putative TATA motifs within 1Kb of CRE | putative TATA motif sequence (5' to 3') | Strand |
|---|---|---|---|---|---|
| **TDRD5 "tejas"_GBI_13502_upstream10Kb** | TGACGTAA | 19 | 2 | VTATAWAWRVVNNNN | plus/ plus |
| **TDRD7 "tapas"_GBI_03370_upstream10Kb** | TGACGTCA | 18 | 4 | VTATAWAWRVVNNNN | minus/ minus/ minus/ plus |
| | TGACGTCA | | 4 | VTATAWAWRVVNNNN | minus/ minus/ minus/ plus |
| **uncharacterized_GBI_15344_upstream10Kb** | TGACGCCA | 17 | 3 | VTATAWAWRVVNNNN | minus/ minus/ plus |
| | TGAGGTCA | | 3 | VTATAWAWRVVNNNN | minus/ minus/ plus |
| | TGACGTCG | | 3 | VTATAWAWRVVNNNN | plus/ minus/ plus |
| **Limkain b1_GBI_15604_upstream10Kb** | TGACGTCA | 10 | 1 | VTATAWAWRVVNNNN | plus |
| | TGACGTCA | | 1 | VTATAWAWRVVNNNN | plus |
| | TGACGTCG | | 4 | VTATAWAWRVVNNNN | minus/ plus/ minus/ minus |

| | | | | | |
|---|---|---|---|---|---|
| **oskar_GBI_01840** | TGACGCCA | 36 | 4 | VTATAWAWRVVNNNN | minus/ plus/ minus/ plus |
| | TGACGTAA | | 3 | VTATAWAWRVVNNNN | plus/ minus/ plus |
| **piwi_GBI_17641** | TGACGTAA | 23 | 2 | VTATAWAWRVVNNNN | minus/ plus |
| | TGACGTAA | | 1 | VTATAWAWRVVNNNN | plus |

| **Sequence name** | **Start** | **End** | **p-value (p<0.001)** | **q-value** | **Matched TATA sequence (5' to 3')** |
|---|---|---|---|---|---|
| **TDRD5 "tejas"_GBI_13502_upstream10Kb** | 3064/ 4668 | 3078/ 4682 | 0.000654/ 0.000843 | 0.8/ 0.823 | TTATTATAAGAATGGC / GTATGAAGACGCCGA |
| **TDRD7 "tapas"_GBI_03370_upstream10Kb** | 8717/ 8092/ 8068/ 7763 | 8731/ 8106/ 8082/ 7777 | 0.000481/ 0.000601/ 0.000638/ 0.00084 | 0.814/ 0.831/ 0.831/ 0.836 | CCATAAATCCCCCTT/ GTATAATGGGGCGGA/ CGATAAAATGGAGTG/ CTATACAAAGAGTCC |
| | 8717/ 8092/ 8068/ 7763 | 8731/ 8106/ 8082/ 7777 | 0.000481/ 0.000601/ 0.000638/ 0.00084 | 0.814/ 0.831/ 0.831/ 0.836 | CCATAAATCCCCCTT/ GTATAATGGGGCGGA/ CGATAAAATGGAGTG/ CTATACAAAGAGTCC |
| **uncharacterized_GBI_15344_upstream10Kb** | 4777/ 6833/ 4691 | 4791/ 6847/ 4705 | 0.00021/ 0.000773/ 0.000981 | 0.725/ 0.916/ 1 | GTAAAAAAGGGGACG/ GTACAAAAACTCCGT/ GTATAGAAGTGAAGG |
| | 4777/ 6833/ 4691 | 4791/ 6847/ 4705 | 0.00021/ 0.000773/ 0.000981 | 0.725/ 0.916/ 1 | GTAAAAAAGGGGACG/ GTACAAAAACTCCGT/ GTATAGAAGTGAAGG |
| | 7647/ 6833/ 8145 | 7661/ 6847/ 8159 | 0.000135/ 0.000773/ 0.000795 | 0.725/ 0.916/ 0.916/ | GTATAAAAAGTAATC/ GTACAAAAACTCCGT/ CTTTAAAAAAAAACGT |
| **Limkain b1_GBI_15604_upstream10Kb** | 5415 | 5429 | 0.000681 | 1 | TTATAAAAGAGAATA |
| | 5415 | 5429 | 0.000681 | 1 | TTATAAAAGAGAATA |
| | 3559/ 2546/ 2867/ 2865 | 3573/ 2560/ 2881/ 2879 | 0.000158/ 0.000594/ 0.00082/ 0.000866 | 1/1/1/1 | GTATAAAAAACAAAT/ CTATAAAATGAATTT/ ATATATAAATATGTG/ ATATAAATATGTGAA |
| **oskar_GBI_01840** | 8725/ 7245/ 8988/ 7026 | 8739/ 7259/ 9002/ 7040 | 0.000499/ 0.000786/ 0.000135/ 0.000293 | 1/ 1/ 0.879/ 1 | GTATTAAAAACAGCT/ CTATTAATGCAAATG/ CTATATAAACAGAA/ TTATAAAGGGGGCAA |
| | 6465/ 6299/ 7026 | 6479/ 6313/ 7040 | 0.000408/ 0.000846/ 0.000293 | 1/1/1/ | ACATAAAAGTCCCTC/ ACATAAAAAAGCACT/ TTATAAAGGGGGCAA |

| | | | | | |
|---|---|---|---|---|---|
| **piwi_GBI_17641** | 2338/ 2832 | 2352/ 2846 | 0.000563/ 0.000308 | 0.688/ 0.688 | CTTTAAAAATGCACA/ GTATAAAAATGTAAA |
| | 5351 | 5365 | 0.000223 | 0.688 | TTATAAAATGGCACC |

**Table S9.** Prediction of putative CRE sites in LOTUS domain-containing *G. bimaculatus* genes. For predictions, p-value was set to less than/equal to 0.0001. The p-value of a motif occurrence is defined as the probability of a random sequence of the same length as the motif matching that position of the sequence with as good or a better score. The score for the match of a position in a sequence to a motif is computed by summing the appropriate entries from each column of the position-dependent scoring matrix that represents the motif. The q-value of a motif occurrence is defined as the false discovery rate if the occurrence is accepted as significant. If there are multiple CRE predictions for one sequence, the table is sorted by increasing p-value for those CRE predictions.

**Supplementary File 1:** The Frequency Matrices (PFM) from JASPAR database used to predict CRE sites and TATA boxes in the presumptive regulatory regions of *G. bimaculatus* genes.

```
TATA box
MEME version 4
ALPHABET= ACGT
strands: + -
Background letter frequencies
A 0.25 C 0.25 G 0.25 T 0.25

MOTIF POL012.1 TATA-Box
letter-probability matrix: alength= 4 w= 15 nsites= 389 E= 0
 0.156812  0.372751  0.390746  0.079692
 0.041131  0.118252  0.046272  0.794344
 0.904884  0.000000  0.005141  0.089974
 0.007712  0.025707  0.005141  0.961440
 0.910026  0.000000  0.012853  0.077121
 0.688946  0.000000  0.000000  0.311054
 0.925450  0.007712  0.025707  0.015424
 0.570694  0.005141  0.113111  0.311054
 0.398458  0.113111  0.403599  0.084833
 0.143959  0.347044  0.385604  0.123393
 0.213368  0.377892  0.329049  0.079692
 0.210797  0.326478  0.329049  0.133676
 0.210797  0.303342  0.329049  0.156812
 0.174807  0.275064  0.357326  0.192802
 0.197943  0.259640  0.359897  0.182519
URL http://jaspar.genereg.net/matrix/POL012.1


CRE consensus sequence full site
MEME version 4
ALPHABET= ACGT
strands: + -
Background letter frequencies
A 0.25 C 0.25 G 0.25 T 0.25

MOTIF MA0018.2 CREB1
letter-probability matrix: alength= 4 w= 8 nsites= 11 E= 0
 0.000000  0.090909  0.090909  0.818182
 0.000000  0.090909  0.909091  0.000000
 1.000000  0.000000  0.000000  0.000000
 0.000000  0.818182  0.181818  0.000000
 0.090909  0.000000  0.909091  0.000000
 0.000000  0.272727  0.000000  0.727273
 0.181818  0.636364  0.090909  0.090909
 0.727273  0.000000  0.090909  0.181818
```

URL http://jaspar.genereg.net/matrix/MA0018.2

**Supplementary File 2:** FASTA files containing the simulated one thousand 10-Kb long DNA fragments generated to test the frequency of occurrence of CRE sites in randomly generated sequences, and LOTUS domain-containing genes.

Download Link
